# Biomimetic model of corticostriatal micro-assemblies discovers new neural code

**DOI:** 10.1101/2023.11.06.565902

**Authors:** Anand Pathak, Scott L. Brincat, Haris Organtzidis, Helmut H. Strey, Sageanne Senneff, Evan G. Antzoulatos, Lilianne R. Mujica-Parodi, Earl K. Miller, Richard Granger

## Abstract

Although computational models have deepened our understanding of neuroscience, it is still highly challenging to link actual low-level physiological activity (spiking, field potentials) and biochemistry (transmitters and receptors) with high-level cognitive abilities (decision-making, working memory) nor with corresponding disorders. We introduce an anatomically-organized multi-scale model directly generating simulated physiology from which extended neural and cognitive phenomena emerge. The model produces spiking, fields, phase synchronies, and synaptic change, directly generating working memory, decisions, and categorization, all of which were then validated on extensive experimental macaque data from which the model received zero prior training of any kind. Moreover, the simulation uncovered a previously unknown neural code specifically predicting upcoming erroneous (“incongruous”) behaviors, also subsequently confirmed in empirical data. The biomimetic model thus directly and predictively links novel decision and reinforcement signals, of computational interest, with novel spiking and field codes, of potential behavioral and clinical relevance.

---

How do collections of neurons somehow produce cognition? How do patterns of physiological activity in cells arranged into anatomical wiring layouts somehow generate perception, learning, memory, decisions, and behaviors? Beyond its critical role in eliciting cognition, the corticostriatal circuit is implicated across a wide range of common yet devastating psychiatric and neurological disorders broadly related to *dopamine, reinforcement learning, conditioning*, and *reward* (1–6). Thus, having a way to simulate the corticostriatal circuit in a biologically realistic manner matters not only because it would answer some of our most fundamental questions in neuroscience, but also because it would permit *in silico* testing of candidate mechanisms for disorders and candidate pharmacological interventions targeting those mechanisms.

In recent years, there have been extraordinary breakthroughs in computational neuroscience. However, most of these models have been optimized for either mechanistic accuracy or to elicit emergent cognition, but not both. On the one hand, there are powerful models with biologically-based systems that reproduce extensive physiologies, from single-cell spiking through generation of oscillations and other critical emergent dynamics. Yet, while these can accurately simulate brain-wide phenomena (7) such as epileptic seizure genesis and propagation (8, 9), they are not designed to generate cognitive processes, such as learning. On the other hand, some of the most remarkable cognitive models are those grounded in artificial neural networks (10–15). Yet, while this is a class of models that effectively “learn,” they do so without reliance on biologically realistic (*biomimetic*) mechanisms at the cellular scale. A small set of influential models optimize for both (16–18). These examples illustrate how a set of constraints from not only the machinery of anatomy and physiology but also the emergent trial-level behavior can be incorporated and satisfied into a single unified model across multiple scales. However, those models were designed to elicit a circumscribed family of behaviors centered on reinforcement learning and were not constructed in a manner that allows them to be easily extended beyond their original scope. Thus, there remains a need for biomimetic cognitive models that can be expanded to encompass a wide range of behavioral processes.

Of course, *in silico* “discovery” of new mechanisms and treatments is possible only if models can apply their learned knowledge to novel or unexpected inputs. A computational model’s ability to make predictions or draw conclusions about data that fall outside the distribution of its training set is referred to as *out-of-distribution inference*. A standard training approach is to train a model from half the data of a particular kind and then test its ability to predict the other half. A far more rigorous standard of generalizability is to ask whether a model built from one kind of data can predict phenomena of a wholly different kind. These latter types of predictions are said to be *zero-trained*.

Here, we address these challenges by building a zero-trained computational model of the corticostriatal circuit designed to be mechanistically accurate across multiple scales, from spiking neurons to local field potentials. This circuit elicits *category learning*, a complex cognitive process encompassing *sensory encoding, perceptual categorization, reinforcement learning, action selection*, and *working memory*. We then validated the model using macaque (nonhuman primate, NHP) electrophysiology and behavioral data (19, 20). Intending to start with a minimal circuit model that would incorporate the above functions, we started by modeling a cortical region, worked our way through to the striatal complex (MSNs, TANs, SNc), and continued further to complete a corticostriatal loop by including globus pallidus (GPi and GPe) and thalamus. Each component consists of a specific assembly of simulated neurons arranged according to anatomical circuit layouts. Each of these micro-assemblies had specific internal structure and anatomical connections, and specific physiological activity characteristics, each as informed by extensive experimental literature, and each such micro-assembly contributed a specific neural computation (*biomimetic computational primitives*, BCPs). Crucially, these neural computations were not assigned top-down but rather emerged from the individual circuit simulations. While we initially expected the corticostriatal circuit to exhibit only a few specific outputs, such as accurate behavior and spectral properties of neural activity, we surprisingly found that the model exhibited far more complex neuronal phenomena than initially intended. Notably, in addition to replicating known physiological patterns and behaviors, the model predicted previously unknown neural encodings, that were then confirmed in the experimental data.

## Results

We developed a generalizable, tractable, biomimetic, multi-scale brain model that categorizes visual stimuli after a delay. The model is zero-trained because it was not directly designed for this task, nor was it “trained” with any empirical data from target datasets, as per modeling approaches based on artificial neural networks. The task for the simulation was chosen following a similar task given to NHPs (macaques) (20), to permit direct comparisons. Simple dot-pattern stimuli (see Methods) were presented to the simulation and to the NHPs, and then, after a delay, the subjects had to decide which of two “categories” the stimulus had been. For NHPs, the decision was signaled by a saccade to the left for one category (A) or to the right for the other (B); the NHPs then either received a reward (for correct category choice) or forced-delay “punishment” (for incorrect choice). The simulation was presented with the same images and delays and produced a corresponding simple simulated left or right motor output, and then received a simulated reward or punishment as in standard reinforcement learning.

Over many “trials” (where the image is shown, a choice is made, and the corresponding reward is given), well-studied plasticity rules from the literature are used to adjust synaptic weights to maximize rewards (see Methods for details). These adjustments lead to improved outcomes over time, and thus learning.

### Details of the brain model at multiple biological and computational scales

The multi-scale model simulates electrical activity of individual neurons, connections among groups of neurons within the same brain region (“assemblies”), and anatomically-based long-distance electrochemical connections between regions. The combination of these interactions, the “connectome,” intrinsically generates the synaptic changes that lead to learning.

There are many ways of describing such a complex model: we provide extensive details and equations in the Methods, Figure 1 shows graphically the hierarchical configuration from individual cortical neurons to the whole brain model, and we describe here the main “subcircuits” that comprise the model. Subcircuits are themselves composed of smaller assemblies of neurons. Many components of the model are contained in more than one subcircuit, so the subcircuits are not defined physiologically, but by specific computational operations.

**Fig. 1.**
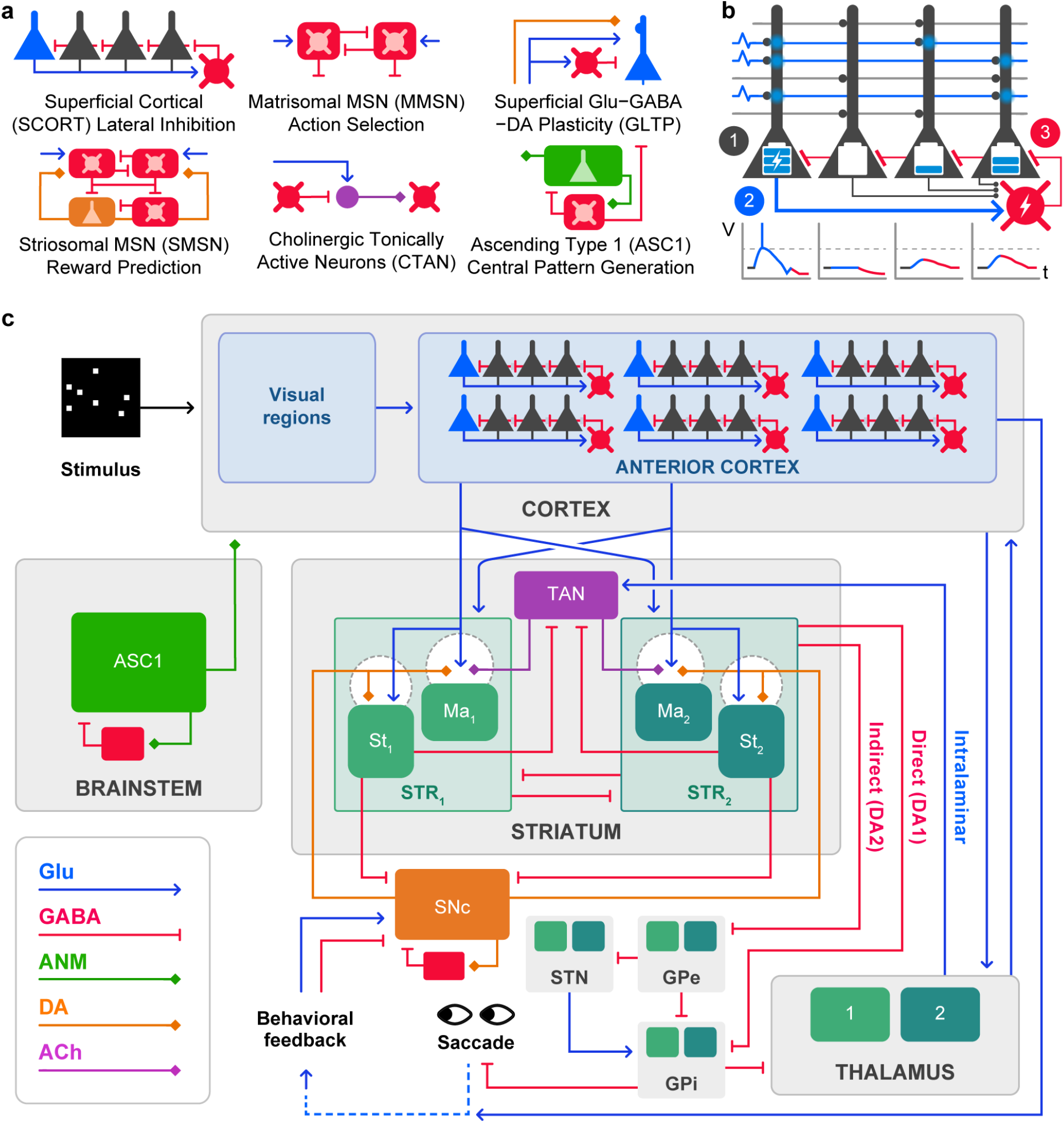
A graphical illustration of our model highlighting key elements and connections from individual neurons organized into assemblies and assemblies organized into brain regions. Detailed discussion in provided in the Methods and the text. (A) Multiple neural assemblies (biomimetic computational primitives), all derived from anatomical structures and physiological neuron operations, and all described in formal detail in Methods; also see Table 1. (B) Closeup on one primitive: Lateral inhibition (see text). This cortical assembly is composed of four excitatory cells and one inhibitory cell. Time-dependent voltage transients within the cells – shown as voltage (V) vs. time (t) graphs – build up over time as a function of the number of active input axons, and the number and weights of connected target dendritic synapses. The overall effect (see text) is that the most-activated excitatory cell(s) reach action potential threshold more rapidly than other cells (1); once any excitatory cell spikes, it activates its local inhibitory cell (2), which in turn inhibits all local excitatory cells (3). Thus, only the most-activated excitatory cell can respond, i.e., the ‘winner’ takes all. (C) Multiple simple cortical ‘columns’ are composed of lateral-inhibition “subcircuits”, which are sparsely connected with each other as described in Methods, as well as receiving diffuse rhythmic input from a simulated simple ascending system (ASC1). Overall model organization, showing how combined cortical and subcortical structures, interconnected as described in the literature, form an integrated simulation with extensive feedforward and feedback elements. The model posterior ‘visual’ regions are presented with visual stimuli, and the system is presented with simple simulated ‘reward’ signals (‘behavioral feedback’) based on the system’s chosen output (saccade) behavior. Cortical structures (visual posterior; prefrontal anterior) are modeled solely as superficial-layer excitatory inhibitory lateral-inhibition assemblies; ascending systems (‘Brainstem’) consist of excitatory-inhibitory oscillatory interactions modeled as next-gen mass models (see Methods). Acetylcholine and dopamine units are modeled from the extensive literature on tonically-active neurons (TANs) and the substantia nigra pars compacta (SNc), respectively. Plasticity rules in cortex and in striatum are modeled separately. Other brain regions are defined as follows: anterior cortex (ANTC), ascending system (ASC1), striatum (STR), striosome (St), matrisome (Ma), globus pallidus pars externa (GPe) and interna (GPi), subthalamic nucleus (STN). Connections among brain regions are each associated with simulated biochemical transmitter / receptor pairs: glutamate (Glu, blue arrows), gamma aminobutyric acid (GABA, capped red lines), ascending neuromodulators (ANM, green lines with diamonds), dopamine (DA, orange lines with diamonds), and acetylcholine (ACh, purple lines with diamonds).

We describe our model computationally by considering the biomimetic computational “primitives” that cannot be further reduced. In computer programming, a “primitive” is a fundamental irreducible language element that performs a simple task (such as adding two numbers). In our model, primitives are realized as assemblies of small numbers of neurons based on specific anatomical layouts of cell types and interactions. The model includes several primitives of physiological computation, such as action selection, reward prediction, central pattern generation, long-term potentiation/reversal, and exploration/exploitation (see Table 1). Some of these are shown in Figure 1A as other biomimetic computational primitives. The composition of primitives produces novel computational operations, allowing for complex behaviors similar to how computer programming primitives can be combined (e.g., in a set of instructions) to perform challenging computations.

**Table 1.** Current Blocks for Corticostriatal Model.

| Biomimetic Computational Primitives | Computational Operations | Regions |
| --- | --- | --- |
| Superficial Cortical (SCORT) | Lateral Inhibition | Anterior, Posterior Neocortex |
| Matrisomal MSN (MMSN) | Action Selection | Caudate, Putamen |
| Striosomal MSN (SMSN) | Reward Prediction | Caudate, Putamen |
| Ascending Type 1 (ASC1) | Central Pattern Generation | Pons, Brainstem |
| Superficial Glu–GABA–DA Plasticity (GLTP) | Long-Term Potentiation, Reversal | Cortex, VTA |
| Cholinergic Tonicly Active Neurons (CTAN) | Exploration/Exploitation | Caudate, Putamen |

For the learning task described here, one important primitive is “lateral inhibition” which takes place within each small assembly of cortical neurons. The anatomical configuration of this computational primitive is an assembly of excitatory and inhibitory cortical neurons shown in Figure 1B. These local assemblies of excitatory and inhibitory neurons in simulated superficial cortical layers are connected such that 1) each glutamatergic input axon (from the simulated visual cortex or any cortico-cortical input) makes contact sparsely with a fixed number of excitatory target dendrites and 2) each excitatory dendrite receives contacts from a separately fixed number of incoming axons. Furthermore, 3) the excitatory cells densely contact local inhibitory cells (in a ratio of about 4:1), and 4) the inhibitory cells reciprocally make dense contacts with local excitatory cells (21–23). Within a local assembly, an activated excitatory neuron activates the inhibitory neuron, which then inhibits all other excitatory neurons (including the one that activated the inhibitation). Thus a very limited subset of target excitatory cells can successfully respond to an input; this “lateral inhibitation” primitive corresponds to the long-known neuronal activity pattern of a specific kind of “winner-takes-all” competitive circuitry. (24–31). Multiple such lateral-inhibition assemblies are present in the cortical structures (visual regions and anterior cortex) in the model, and they are sparsely interconnected with each other, as briefly illustrated in Figure 1C, and discussed in Methods.

The evolution of neuronal activity in these assemblies starts with an incoming set of spikes from a simple “visual” cortex region (SVCR). The stimulus (dots on the screen) activates these neurons, with individual connections corresponding to specific pixels on the screen. These neurons induce depolarization via input axons that contact dendritic synapses of neurons in the simulated anterior cortical area (ANTC) as illustrated in Figure S2. The total voltage over time (illustrated by voltage-time graphs) rises at a rate that is dependent on the quantity of input activation at a given target cell (e.g., the brightness of that pixel) and the weights of the receiving synapses. The active synapses of the responding cells will then be increased slightly in weight as determined by the rules from the literature for glutamatergic synaptic plasticity (see Methods). These synaptic increases makes it even more likely that slight variants of an input will activate the same target excitatory cells. As a result, similar inputs will increasingly come to elicit identical, not just similar, cortical responses (Figures S6-S7).

A set of information-processing operations is thus carried out by a simple, but biologically grounded, small assembly of excitatory and inhibitory cells. This assembly of typically 5-6 neurons is irreducible and can thus be considered to be one functionally ‘primitive’ element of brain circuitry. The arrangement used here is very distinct from ANN approaches to generating lateral inhibition (25–31). It also provides a starting point for investigating how biochemical deviations (e.g., pharmacological blocking or modulation of glutamatergic or GABAergic receptors) could affect cognitive processes.

At a larger scale, our model incorporates well-studied physiological properties of neurons into four subcircuits (Figure 1C). The *Cortex Subcircuit*, comprising the aforementioned Simple Visual Cortical Region (SVCR) and Anterior Cortex (ANTC), is composed of blocks of lateral inhibition assemblies that receive input from a simulated external visual grid input (“Stimulus” in Figure 1C) with some connections between assemblies. This subcircuit also receives ascending rhythmic inputs (ASC1 in Figure 1C) modeled as next-generation neural mass models (32, 33) as described in Methods. The *Striatal Subcircuit* consists of matrix (matrisome) assemblies, patch (striosome) assemblies, and a substantia nigra pars compacta (SNc). Matrisome and striosome blocks are composed of medium spiny neurons (MSNs) receiving convergent inputs from the cortex (via glutamate), from the Tonically Active Neuron (TAN) stucture (via acetocholine), and from the SNc (via dopamine) while outputting to the pallidum and thalamus. The *TAN Subcircuit* is composed of cholinergic TANs that receive inputs from intralaminar thalamic nuclei (34) and GABAergic striosomal MSNs. TAN projection synapses co-localize with corticostriatal synpases. Neurons in both the Striatal and TAN Subcircuits are not modeled as spiking neural elements but as simplified physiologically-derived “rules” whose equations define their input/output behavior (see Methods). The *Brainstem Subcircuit* is a highly simplified ascending system (ASC1 in Figure 1 C). This represents a simulated ascending system (such as locus coeruleus or basal forebrain) that generates oscillatory input, centered around 16 Hz. The resulting rhythmic activity provides input to the ANTC beyond the SVCR input generated by the stimulus. It is modeled using next-generation mean-field neural mass models (32, 33) and descending signals are not considered. Details on these subcircuits are provided in the Methods and Supplementary Materials.

The interconnections between these subcircuits support the process of learning. As the model experiences many trials, there are changes to synaptic strength for cortico-cortical, cortico-striatal, and thalamo-cortical contacts that cause changes in behavior. For example, initial trials have significant cholinergic “noise” from the TAN Subcircuit that increases variance in target matrix MSN responses, which we hypothesize adds “exploratory” variability to striatal action selection (see 35). As learning progresses, corticostriatal synapses to striosomes are strengthened, which increases their ability to inhibit TANs, which decreases variability as associations are effectively “learned” in the corticostriatal system. Thus, the TAN subcircuit is a crucial part of an “exploration/exploitation” primitive operation (see Figure S3).

### The model succeeded in categorical learning at a rate similar to NHPs while replicating cortico-striatal synchrony effects

Our model is intended to characterize phenomena that currently cut across multiple levels of description. For example, in real brains, spiking neurons give rise to waves, yet waves in turn strongly modulate spiking; and both spiking and waves appear affected by disorders and by drugs (e.g., 36–39). We provide initial evidence that the simulation may assist in uncovering the mechanisms of these cross-level relations.

Preliminary validation of the model came from the observation that the simulated neuronal activity and interaction patterns were similar to NHP brains as they perform similar tasks (see Figures S4 and S5). Although the simulation carries out solely physiological operations, in doing so, it successfully learns the visual categorization task that it is presented with. That is, via synaptic change, it acquires the category characteristics (i.e., discriminating category A from B) and it learns (via striatal-based reinforcement) to produce simple appropriate behaviors (a simulated “saccade” left or right).

The behavioral responses increase in accuracy across trials (Figure 2A) in both simulation (top) and the experimental animals (bottom) and at comparable rates. Cortico-cortical synapses exhibit long-term increases in weight (Figure 2B). By contrast, striatal synapses exhibit increases and decreases as per the combined effect of cortical glutamatergic, SNc dopaminergic, and TAN cholinergic inputs that converge at each spine of striatal medium spiny neurons (see Methods and Figure 1). Thus, cortical synapses are seen to slowly strengthen, while corticostriatal synapses strengthen and weaken over trials.

**Fig. 2.**
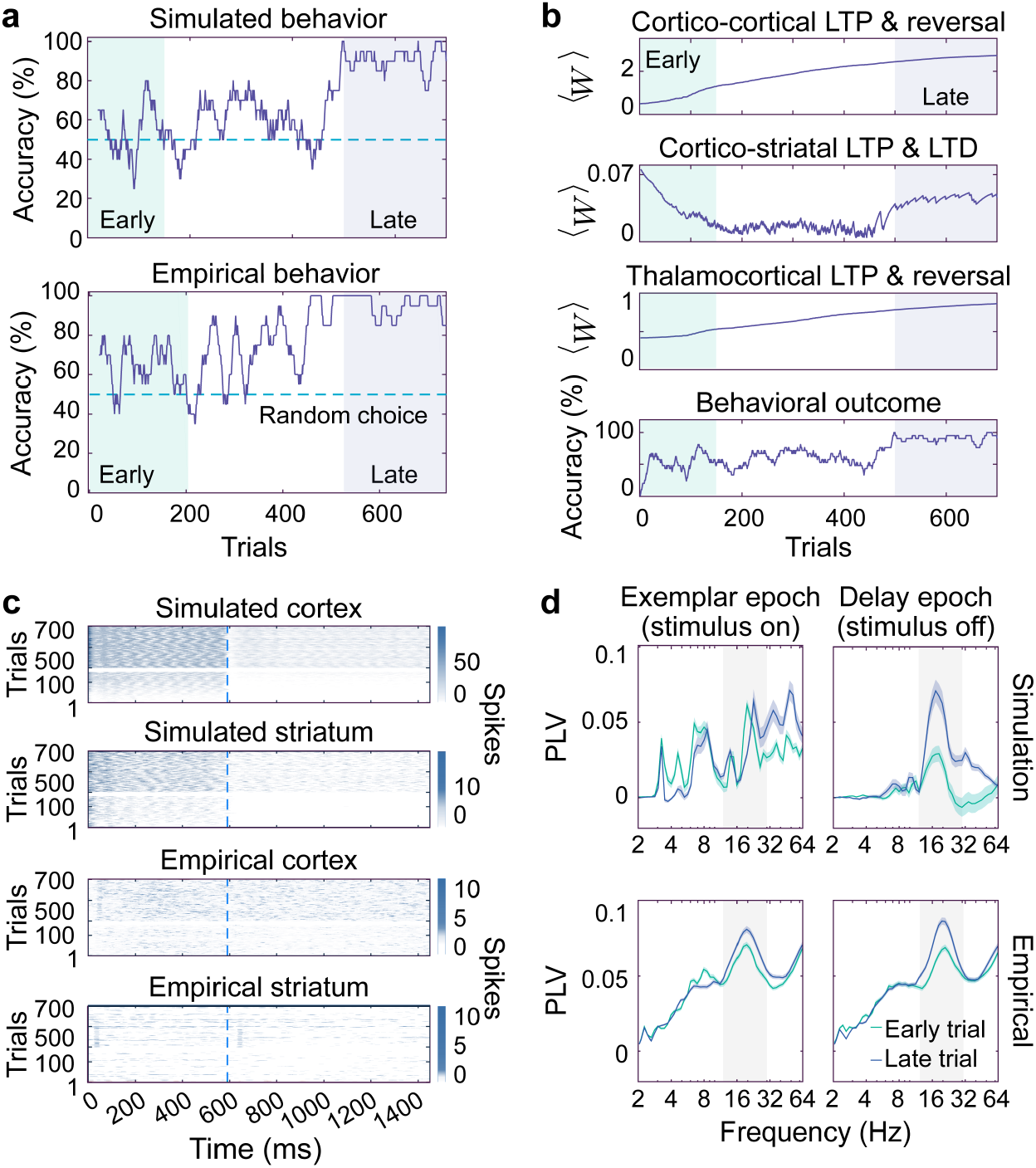
Cortical and striatal synaptic change underlie changes in spiking and synchrony leading to behavioral changes in the form of learning. (A) Behavioral learning over trials in simulation (top) and NHPs (bottom) exhibit comparable characteristics. Correct behaviors increase slowly over early trials (0-200, light green shaded), and reach plateau performance by about trial 300. We define trials 500-700 as “late” (gray shaded). (B) Changes to simulated synaptic strength for cortico-cortical, cortico-striatal, and thalamo-cortical contacts across trials, as well as accuracy of behavioral trials. The trends show Long Term Potentiation (LTP), Long Term Depression (LTD), and synaptic potentiation reversal. (C) Simulated (top two) and empirical (bottom two) activity of single sessions across trials. Each row represents Local Spike Summations (for simulated data) and Local Field Potentials (for empirical data) over a single trial. Dashed lines indicate the time of stimulus offset. (D) Corticostriatal phase-locking values (PLV) from simulated (top) and empirical (bottom) data. These show that later trials (gray) had increased phase locking between the cortex and striatum in the beta frequency band (in yellow) which is driven by the simulated ASC1. This phase locking remains even when the stimulus is removed for both the simulation and the NHPs.

The process of learning and of holding information in “working memory” depends on interconnections among all four primary (cortical, striatal, TAN, and brainstem) Subcircuits. Ongoing recurrent activity, via feedback from striatum to thalamus to cortex to striatum, drives the ongoing memory, rather than any specific specialized cell states in striatal or anterior cortical cells. This is concordant with the recorded data from the NHPs, but it is not yet known whether more detailed representations of striatal or anterior cortical cells would change these activities.

Investigating neuronal activity differences between early and late trials (Figure 2C) shows that spiking is much stronger in late trials for both simulated and empirical systems, in both the cortex and striatum, and both during and after stimulus presentation. Empirical observations of spiking in NHP brains are quantified using the “Local Field Potential” (LFP) (20). For the simulation, we developed a similar metric called Local Spike Summation (LSS), a simple measure of simulated neuronal activity defined in Methods. While there are many similarities in these measures of spiking between the empirical and simulated brains, one noticable difference is that NHP brains do not show as marked a decrease in spiking after stimulus offset as the model (Figure 2C). Of additional interested is the “synchrony” between cortical and striatal activity which is calculated using the “Phase Locking Value” (PLV) from a wavelet analysis of LSS or LFP time series (see Methods and Supplementary Materials) as was done in (20). PLV is measured in multiple frequency bands including the near 16 Hz (beta-frequency) band driven by the ASC1 in the simulations. Figure 2D shows that both simulated and empirical data showed increased synchrony in the beta-frequency band in late trials versus early trials, though this effect was much stronger in the simulation.

Comparison of activity during correct and incorrect trials indicates that synchrony in the beta frequency band is limited to correct trials and caused by changes in weights. To compare correct trials vs. incorrect trials, the difference in corticostriatal synchrony (measured by PLV) is shown in Figure 3. Both simulated and empirical results show that slightly higher synchrony during early incorrect trials is reversed with learning so that later trials show stronger synchrony in the beta frequency band only on correct trials (Figure 3A). The simulation provides a candidate explanation for the increased synchrony in the form of higher cortico-striatal synaptic weights for neurons that respond on correct trails. The weights for correct and incorrect neurons are similar for about 450 trials and then markedly diverge as weights for “correct” neurons double (Figure 3B). These weight differences for correct and incorrect trials are also reflected in the rate of neuron spiking which is much lower and more localized in both the cortex and striatum during incorrect trials (Figure 3C).

**Fig. 3.**
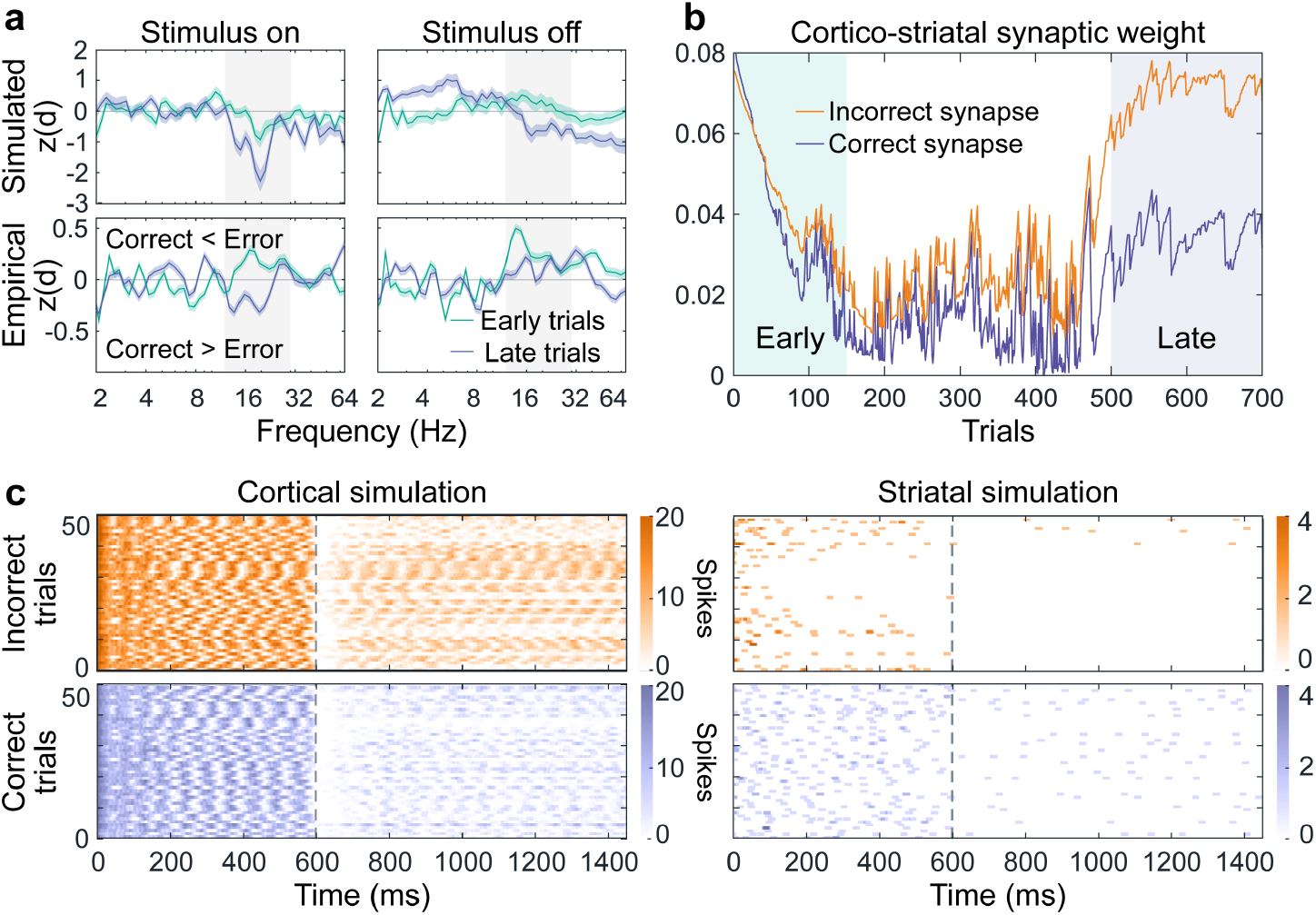
Strengthening of cortical-striatal synaptic weights support phase locking values and making correct choices. (A) Corticostriatal synchrony on correct vs. incorrect trials as measured by Phase Locking Values (PLV) at different frequencies (with the beta frequencies driven by the simulated ASC1 highlighted in yellow). Shown are PLVs for incorrect minus correct trials on average, in units of standard deviations from a random sample (e.g., they are z-transformed, see Methods). Values above zero are instances where incorrect trials have greater synchrony than correct trials; less than zero means correct trials have greater synchrony. In the model (“simulated”, top), corticostriatal synchrony during incorrect trials in the beta frequency range decreases with learning, from early (light red) to late (dark red) trials, both during stimulus presentation (left) and delay period (right); the macaque subjects (“empirical”, bottom) exhibited this same effect as reported in (20). (B) In the simulation, the synchrony decrease is found to be due predominantly to differential synaptic strengthening on those striatal cells responding on correct (blue) vs on incorrect trials (red). (C) These synaptic effects show up in local spike summation (LSS) plots of simulation population activity in late trials, after extensive learning, in cortex (left) and striatum (right) on incorrect (top) and correct (bottom) trials. Due to selective weakening of corticostriatal synapses on incorrectly-responding cells (as per panel B), the striatal responses in incorrect trials become extraordinarily sparse; this is the primary contributor to diminishing corticostriatal phase-locking values.

### The model has predictive power: the discovery of incongruent neurons

An effective model of the brain will provide significant new insights into neuroscientific questions. Here we show that our model has specific predictive power as illustrated by the discovery of “incongruent neurons” in the simulation. Although we report in parallel the discovery in simulated and empirical data, we emphasize that the simulation results prompted the successful identification of a previously-missed behavior in the long-studied empirical data.

Analysis of the simulation showed surprising predictive behavior in some neurons. After extended learning trials in the category task, our simplified brain model reached a phase where every simulated excitatory anterior cortical neuron had a characteristic preference to fire for stimulus category A or B. Unexpectedly, every such neuron also had a specific response to the *outcome* of the trial: correct vs. incorrect. That is, all simulated neurons were found to show strongly selective responses to one of four conditions: A-correct, A-incorrect, B-correct, and B-incorrect. These selective responses were visible after only 200 ms (see Figures S4, S5, and S8). An initial expectation would be that a “trained” model that had successfully learned to distinguish between the two stimuli would have optimized its performance by having practically all neurons supporting correct choices. Those neurons that strongly correlated with an incorrect choice we thus defined as “incongruent neurons.” These incongruent neurons (ICNs) learned to predict an upcoming behavioral error. Commprising about 20% of all neurons, ICNs sufficiently activate target striatal action-selection cells for the “other” category, thus causing the wrong behavioral response. To explore these in greater detail and to ascertain evidence for incongruent neurons in the empirical data, we use a simple method to identify and characterize incongruent neurons. 1) Trials are sorted into the four classes (A-correct, A-incorrect, B-correct, B-incorrect). 2) Spiking rates of each excitatory anterior cortical neuron during the first 200 msec of a given trial were normalized by the average number of spikes across the population for that trial-type. This then allows for 3) assignment of a neuron to one of the four classes based on the highest spiking rate (“activity”). This simple sorting by their predicted decisions leads to neurons strongly (and often uniquely) correlated with each of the four classes as shown in Figure 4A. Though the macaque implanted electrodes acquire recordings of only very few neurons, Figure 4A does indicate that the empirically recorded neurons appear also to exhibit predictiveness of specific eventual behavioral outcomes.

**Fig. 4.**
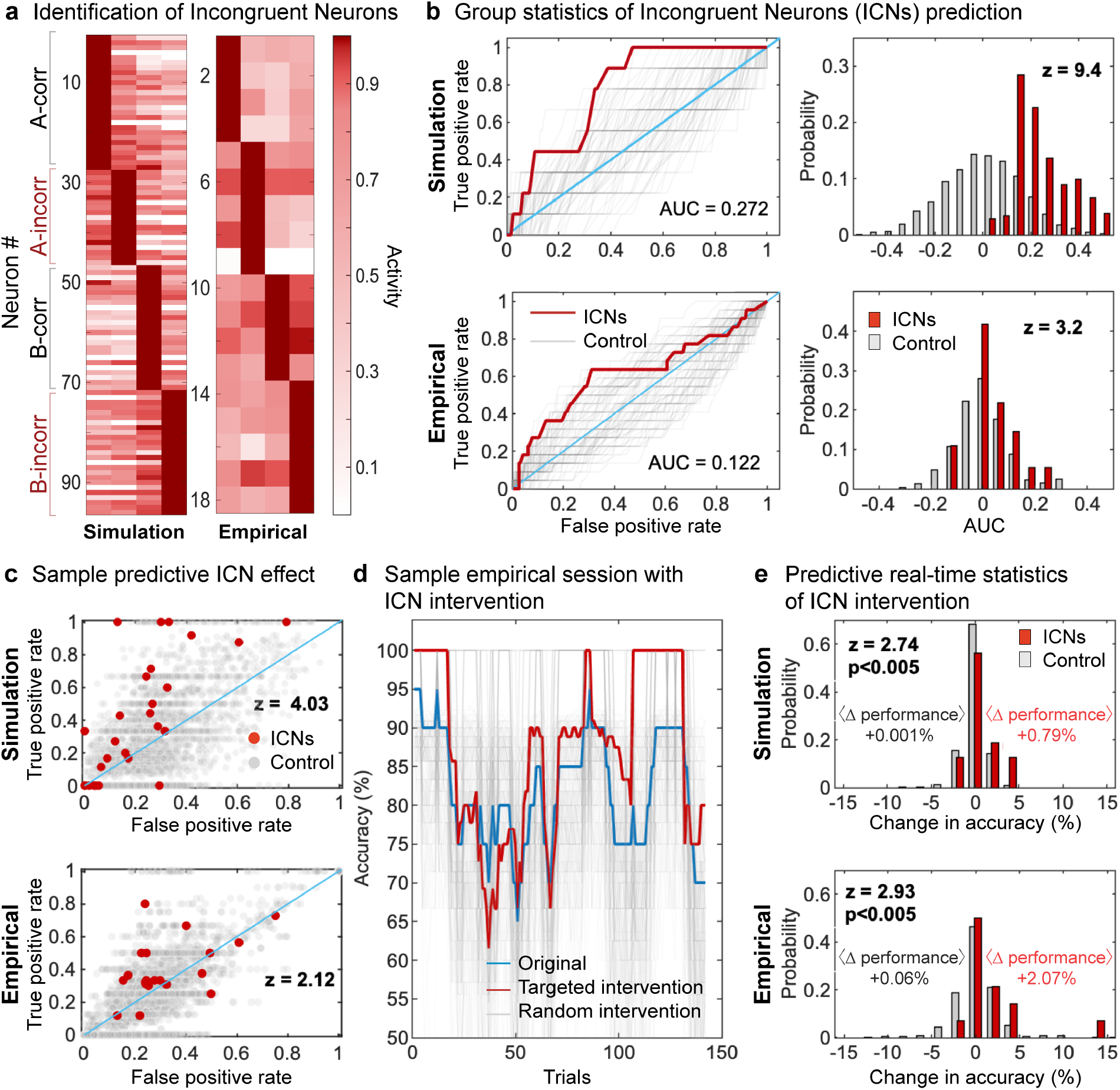
“Incongruent Neurons” (ICNs) were discovered in the simulation and later confirmed in empirical data: they reliably predict incorrect choices long before they occur, and are statistically highly significant. (A) Simulated and empirical cortical spiking preferences (see text) sorted by four conditions over the first 200 msec. (B) ROC (Receiver Operating Characteristic) plots showing how True Positives exceed False Positives for ICNs as measured by area under the curve (AUC). ICNs significantly outperform randomly chosen neurons, showing that the categorization into congruent and incongruent neurons is meaningful. (C) ROC for ICNs per session using a predetermined fixed decision threshold (see Methods); red dots are ICN predictiveness and gray dots are predictiveness of randomly chosen cells. (D) Sample trial-by-trial run showing effects of a real-time predictive intervention that halts trials based on sensing of bad-idea neuron spiking patterns (red) or random spiking (gray) compared to the original performance (blue). (E) Summary statistics over all sessions: random neurons (gray) have negligible ability to change accuracy of outcomes (0.06±0.72%) in empirical trials whereas ICNs (red) increase accuracy significantly (2.07% for empirical NHP trials and 0.79% for simulated trials). This level of improvement by random chance happens with probability *p*<0.005. The increased accuracy confirms that NHP brains would improve at the task if ICNs responded differently and that ICNs are much better at predicted behavioral errors than random neurons.

We confirmed that ICNs were self-consistent and persistent across late trials in both simulated and empirical data with strong statistical significance. To ascertain statistical validity, we divided the late trials in half, and used the first half for determination of purported ICNs which then were tested by being applied to the separate second half of late trials. Figure 4B shows that ICNs predict behavioral outcomes very significantly better than randomly chosen neurons (*p <* 0.001 for empirical data and *p <* 10^−21^ for simulated data).

We then applied an even more stringent threshold for validation of ICNs: *could these neurons predict outcomes, in real-time, within individual trials?* If a hypothetical device were implanted into the NHPs and trained on initial trials, it could be tested for its ability to improve behavior, by halting any trial in real-time whenever incongruent neurons predicted an upcoming erroneous response. Figure 4D shows a sample trial-by-trial run where such an ICN-based intervention significantly (p<0.005) increases behavioral accuracy (Figure 4E) of the simulation.

Finally, we identified specific aspects of the simulation that show how ICNs are mechanistically produced. Supplementary Figure S10 begins with sparse synaptic cortical connectivity and maps those connections backward, through multiple cortical areas, superimposing them directly onto the visual input stimuli. The sparse combination of synapses that eventually co-occur at downstream cortical cells turns out to define circumscribed visual receptive fields that any given cortical neuron can “see”. Analyses (in Supplementary Figure S10 B-D) show that cortical cells are thus anatomically “pre-weighted” to selectively respond probabilistically either to category A or category B stimuli. These anatomical biases reliably predict eventual learned prefrontal category A vs B cells, whereas less-biased, or more-ambiguous cells, become candidate incongruent neurons (ICNs). These detailed analysis of anatomical bias and synaptic modification are further detailed in Supplementary Information section IV.

These results statistically validate that ICNs are a real feature of both simulated and, surprisingly, NHP brains. Thus, our corticostriatal circuit demonstrates its ability to satisfy the strictest bar required of zero-trained models: providing proof of principle that it can be used to “discover” new phenomena qualitatively distinct from the information used to encode it.

## Discussion

We demonstrate a brain circuit model capable of simulating activity on multiple scales, from individual neurons, to anatomical micro-assemblies, to whole brain rhythmic interactions, to behavior. The model is built solely from mechanistically accurate neural microcircuits, each corresponding to distinct brain region structures, and each of which naturally elicits distinct computational processes. These processes are shown to contribute basic operations, such as lateral inhibition and exploration/exploitation, to a complex categorical learning task. Model outputs reliably match independent experimental data acquired from NHPs, even though the model received zero training from any of these data. The model’s behavior includes detailed spiking behavior and cortical-striatal synchrony; learning-related changes to these spiking and field responses across trials; the accuracy of the task as a function of the number of trials; and selectivity of individual cortical neurons both for visual stimulus category and for trial outcome (correct vs. incorrect); all of these are shown to correspond to data from NHPs performing an equivalent series of tasks.

Other models such as *Humphries-Stewart-Gurney (HSG)* (16) and *Ashby-Crossley-Horvitz (ACH)* (17, 18) share our goal of adopting constraints from physiology in their accounts of high-level cognitive abilities. HSG presents a mechanistic model that incorporates detailed anatomical and physiological corticostriatal data to demonstrate how the basal ganglia can perform action selection and how oscillatory activity can emerge from interactions between the constituent nuclei. ACH addresses automaticity in perceptual categorization via reinforcement learning arising from the basal ganglia. Their model also investigates how cholinergic interneurons in the striatum can protect striatal-dependent learning from interference, allowing for the acquisition of new skills without overwriting previously learned ones, demonstrating the importance of incorporating real biological constraints into such models.

Our approach fundamentally differs from these efforts by utilizing a modular architecture based on BCPs; these biologically informed computational structures enable significant augmentation of the model’s scope, encompassing a broader range of cognitive operations while providing building blocks for further expansion. Primitives as used in computational neuroscience (40, 41) are “inspired by”, or can be more directly derived from, the properties and functions of biological neurons and neural circuits, proposing abstractions of complex neural processes into mathematically tractable components. Examples include *neural activation functions*, such as sigmoids and rectified linear units (ReLU) (40, 41), *synaptic plasticity rules*, like Hebbian learning and spike-timing dependent plasticity (STDP) (42, 43), *neural network architectures*, including feedforward networks and recurrent neural networks RNNs) (44, 45), *probabilistic inference*, such as Bayesian inference and Markov decision processes (MDPs) (46, 47), and *attractor dynamics*, like Hopfield networks and continuous attractor neural network (CANNs) (48, 49).

By combining these primitives in various ways, computational neuroscientists can develop testable hypotheses and gain insights into the underlying mechanisms of neural computation, from single neuron dynamics to large-scale network interactions (40, 41). Computational primitives can be phenomenologically predictive without being mechanistically biomimetic. A recurring open question is how predictive synthetic primitives can be across wide realms of distinct neural and behavioral data, instead of models that may individually predict certain experiments but must be changed to correspond to further data. One of the most commonly used synthetic computational primitives in computational neuroscience is *back-propagation*, or *backprop*, a supervised learning algorithm that trains artificial neural networks, particularly multi-layer perceptrons (MLPs) (50) such as “deep learning” networks (44), using gradient descent to minimize the difference between the network’s output and a “desired output” that must be separately provided to the network. Confusingly, the terms *computational primitives* and *motifs* are sometimes used interchangeably, to variously mean fundamental operations within computational models (40, 41), or to recurring structural patterns of connectivity or dynamics such as feed-forward loops, recurrent loops, and central pattern generators (51, 52). To clarify, what we term BCPs describe *physiological computation*: bottom-up mechanistic information-processing operations that arise directly from the physiological steps taken by cells within well-defined anatomical circuitry from distinct brain regions. We specifically posit that these operations combine to give rise to brain-based cognition, thus enabling the prediction of both information-processing and behavioral operations as well as the specific electrophysiological and biochemical mechanisms that directly underlie those operations.

One of the most powerful implications of this BCP-based model is that its simulation identified an entirely novel class of neural signals, which were then independently verified using NHP data. Termed “incongruent neurons” (ICNs), these cells’ early activity in the task predict an upcoming incorrect choice that occurs far later. Notably, ICNs correspond with very large local field potential (LFP) effects that show up prominently in NHP recordings of cortico-striatal synchrony (see Figure 3). Therefore, ICNs appear to be integral to a large-scale signaling system coordinating activity between the cortex and striatum. This predictive activity of ICNs is consistent with other contexts in which neural activity patterns in the brain were found to predict later motor “decisions” (53–57), yet also are markedly distinct in that they predict error. ICNs predict an undesirable action that the animal is about to carry out; the animal is “choosing” to do something presumably against its own interest. What functions might ICNs play in predicting error? The NHP experimental subjects were not constructed merely to solve only this particular two-category task, nor was the simulation. Rather, the simulation was solely based on physiological computation as described. Quite different sets of environmental challenges may be presented (to the NHP or the simulation), containing multiple categories (and subcategories), establishing entirely distinct pairings of input types and response types. In this wider world, ICNs may play a functional role in avoiding local minima, adding flexibility or alternate choices to the organism’s decision-making mechanisms. Ongoing work in reinforcement learning continues to address central questions of how predictive and retrospective information is neurally coded (58); the discovery of neural ICN coding may help our understanding. This incongruence among decision, intention, and valence offers a rich panoply of directions for future study.

Finally, we emphasize the broader point that it was the zero-trained nature of simulations based on biomimetic computation that enabled the emergence of these broad and unexpected findings. Whereas much computational work focuses on the statistical learning of specific data, the present work instead attempts to uncover how low-level physiological spiking, embedded in specific delimited sets of anatomical wiring organizations, can itself give rise to the forms of neural and behavioral results that brains produce. The simulation does not learn from any specific data, but rather is constructed from a large cited corpus of very well-known and highly cited anatomy and physiology. As a result, the simulation does not just learn in a way that is specific to a particular set of stimuli or demands but rather generates a broad range of functions that encompass the phenomenological realms of sensory encoding, perceptual categorization, reinforcement learning, decision-making, action selection, working memory, and category learning, as well as extensive neural patterns of spiking, field potentials, synchrony, phase-locking, and changes to these physiological patterns that occur with synaptic change. The breadth and depth of the findings from this zero-trained model are powerful evidence of its utility in the analysis of existing data, and are suggestive of its potential applicability to a broad range of challenging questions from the neurobiological bases of behaviors and associated disorders linked to predictive coding and reinforcement learning.

## Methods

We describe the corticostriatal circuit from mechanistic to emergent scales. We begin with a detailed model description at the neuronal and assembly levels. We then provide details on how corticostriatal decision-making proceeds from synaptic plasticity, e.g., how the neurons are continually modified to proceed with learning. We then provide details for our computational methods, the experimental task design, the simulation protocol, and how we analyze the model’s physiological response. See the Supplementary Information Methods section for additional information regarding the model’s construction, parameter values, and references.

### Detailed model description

#### Extensive anatomical and physiological features incorporated into the present simulation

A sizeable body of references was used in the conception and construction of the several subcircuits and neural assemblies that constitute the present brain circuit simulation (Table 1, Figure 1D); these consist of attributes that are highly cited in the literature. For list of key publications for each computational primitive, see Table A1 in Supplementary Materials.

### Spiking neurons and synaptic connections

#### Neuronal Dynamics

Each neuron is modeled using ionic conductance based point neuron models following the Hodgkin-Huxley formalism (59). For simplicity, in this model we consider only voltage gated Sodium and Potassium channels. The membrane potential is described by:

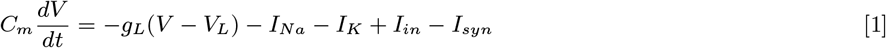

where *I*_*in*_ is the input current, which can either be external stimulus or background activity, and *I*_*syn*_ is the synaptic input. Sodium current 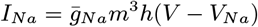 and potassium current 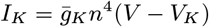 are driven by gating variables {*m, h, n*} whose dynamics are governed by the general form :

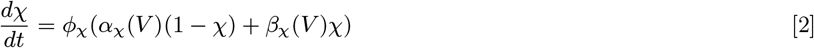

Here, *χ* is the gating variable and *ϕ*_*χ*_ is the temperature related effect on the time scale. See Methods section in Supplementary Information for parameter values.

### Synapse

Synaptic current in Eq. 1 for a neuron *i* is the sum of synaptic inputs from all incoming axons {*j*} viz. 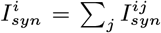 Synaptic current from neuron *j* to neuron *i* is given by

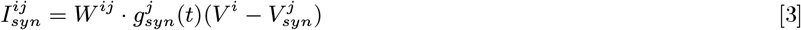

where *W* ^*ij*^ is the synaptic weight from neuron *j* to *i*, a quantity which estimates density of dendritic spines and is modified by Long Term Potentiation (LTP) and Long Term Depression (LTD, see Eq. 28,30). The synaptic conductance 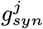 is a function of spiking activity of the presynaptic neuron, and therefore is one of the state variables for each neuron whose dynamics is described by damped oscillator equation:

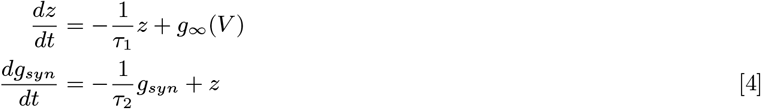

which is driven by impulse force *g*_∞_(*V*) defined as follows (60)

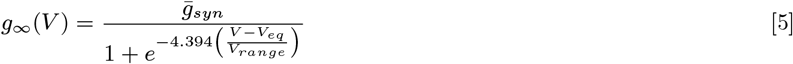

See Methods section in Supplementary Information for parameter values.

### Brain regions modeled as cell assemblies

The schematic of the overall circuit is shown in Figure 1. Most of the brain areas shown in the circuit have been modeled as cell assemblies of sparsely connected spiking neurons, where each region has its own specific internal structure and neuronal types comprising them. These structures and neuronal types are derived from the well known anatomical and physiological literature. Below are the specific structures these brain areas modeled here.

### Cortical areas

Superficial layers of cortical regions in this model (cortical blocks) comprise large number of local lateral-inhibition circuits (winner-take-all). Each winner-take-all circuit (see Fig. 1 A), comprises 5 excitatory neurons sending synapses to one feedback inhibitory neuron while the inhibitory neuron sends synapses back to the 5 excitatory neurons. The winner-take-all circuits are connected to each other through synapses between the excitatory neurons. We have set the connection density such that each excitatory neuron is connected to 1 excitatory neuron from the same cortical block. Within each cortical block, we have 1 *feed-forward* inhibitory neuron which recieves input from excitatory neurons of other cortical regions and sends inhibitory synapses to excitatory neurons of the same cortical block. These feed-forward neurons also receive modulatory inputs from various *Brainstem* nuclei forming the Ascending Reticular Activating System (61). The rhythmic bursting of feed-forward inhibitory neurons causes the rhythmic bursting of the excitatory neurons and which in turn sets the dominant rhythm in the cortical region.

There are two cortical blocks in this simplified model: the Simple Visual Cortical Region (SVCR) and the *Prefrontal Cortex* (PFC). The SVCR consists of 45 winner-take-all circuits comprising (45 × 5 =) 225 excitatory ‘target’ neurons which receive the stimulus dot patters in the form of input current *I*_*in*_. The visual stimulus consists of (15 × 15 =) 225 pixels (see Figure S6) which are mapped directly onto the 225 visual area neurons, where the *I*_*in*_ for each neuron is proportional to gray scale value of the corresponding pixel.

The Prefrontal cortex (PFC) is comprised of 20 winner-take-all circuits, hence (20 × 5 =) 100 target excitatory cells. The connections between the SVCR and the PFC are hypergeometric (62) viz. every neuron has equal number of outgoing and incoming synaptic connections. The resulting hypergeometric connectome (62) ensures the absence of “hot” or “cold” neurons, i.e., cells differently capable of activity due solely to their afferent wiring. Every SVCR neuron makes an outgoing synapse with 8 out of 100 PFC neurons, while each PFC neuron receives incoming synapses from 18 out of 225 SVCR neurons, thus the connection probability for cortico-cortical connection is 8%.

### Striatum

For the striatal-pallidal projection system, we model two blocks (‘*L*’ and ‘*R*’, say) for each of the downstream brain regions in the loop, viz. *Striatum, GPi, GPe, STN*, and *Thalamus*. Each block in one region connects with the corresponding block in the other region viz. *L*(*R*) block of one region connects to *L*(*R*) block of another region, thus representing the topographic connections. These two streams of connection correspond to the two choices of response actions (in this case left or right saccade).

The Striatum has three components: *Matrisomes* (matrix) and *Striosomes* (patch) Medium Spiny Neurons (MSNs) that are GABAergic, and the *Tonically Active Neurons* (TANs) that are cholinergic. These three form a striatal complex (35), which is supposed to play a vital role in reward-based action selection (63). We modeled these components along with *SNc* as simplified physiologically-derived “rules” defining their input/output behavior (see following subsection, Eq. 11,14,16,18). However, we also modeled two blocks of matrisome as spiking neuronal assemblies (*L-STR* and *R-STR*) to capture the temporal and spectral properties of the striatal data from the experiments. We also model a single TAN spiking neuron which provides modulatory input to MSNs.

Each matrisomal block consists of single layer of 25 inhibitory neurons, where each neuron gets incoming synapses from 4 PFC neurons. These neurons also get sub-threshold excitation from a TAN neuron. Since it was experimentally observed that TAN neurons are phase-locked with the striatal LFPs (64, 65), we propose that the striatal rhythm is modulated by rhythmic bursting of TANs. TAN neurons in turn get the rhythmic input from *Intralaminar Thalamic Nuclei*.

### GPe

Two blocks of single-layer assemblies of inhibitory neurons, 15 neurons in each block. Each neuron is connected to 1 striatal neuron of the corresponding block in striatum (topographic connections). Each GPe neuron is connected to 1 GPi neuron of corresponding block.

### STN

Two blocks of single-layer assemblies of excitatory neurons, 15 neurons in each block. Each neuron is connected to 1 GPe neuron of corresponding block in GPe (topographic connections). Each STN neuron is connected to 1 GPi neuron of corresponding block.

### GPi

Two blocks of single-layer assemblies of inhibitory neurons, 25 neurons in each block. Each neuron is connected to 1 striatal neuron of corresponding block.

### Thalamus

Two blocks of single-layer assemblies of excitatory neurons, 25 neurons in each block. Each neuron is connected to 1 GPi neuron of the corresponding block. Thalamic neurons project back to PFC target neurons thus completing the loop (see Figure 1). Each PFC neuron gets input from 16 thalamic neurons i.e. 8 neurons from each thalamic block.

### Ascending Systems and Intralaminar Thalamic Nuclei

The simulated ascending system (ASC1) is an extreme simplification of ascending systems such as the cholinergic basal forebrain and noradrenergic locus coeruleus; it produces rhythmic activity via a neural mass model. That activity becomes simulated modulatory input to cortex, driving a beta rhythm in the cortex(66). The ASC1 simulation thus functions as a central pattern generator. It has been modeled using a simple ‘Next Generation’ neural mass model (32, 33) which is a mean field approximation of a network of rhythmic neurons. A single neural mass representing an ascending system nucleus comprises a synaptically coupled population of excitatory (E) and inhibitory (I) neurons. Their collective activity is defined by the complex valued Kuramoto order parameter *Z*_*e*_ and *Z*_*i*_ respectively, while their synaptic interaction is measured by synaptic conductivities *g*_*ee*_, *g*_*ei*_, *g*_*ie*_ and *g*_*ii*_ such that *g*_*ab*_ is the synaptic conductivity from population *b* to population *a*, where *a, b* ∈ {*E, I*}. The dynamics is described as :

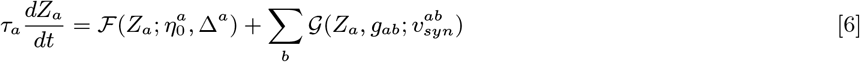

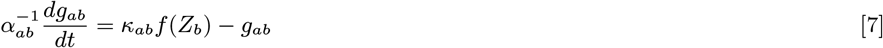

where

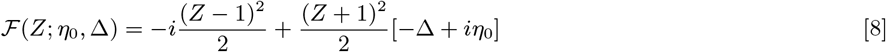

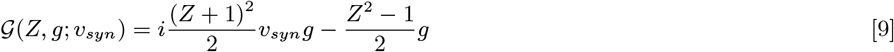

and the population firing rate *f* is given by

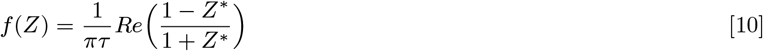

The firing rate *f* (*Z*_*e*_) of population *E* directly affects the modulation of the target neurons, in the form of an input current to the target neurons such that *I*_*mod*_ = *k*_*LC*_*f* (*Z*_*e*_). In this simulation, those targets are simulated somatostatin feed-forward inhibitory cortical neurons. See Methods section in Supplementary Information for parameter values.

### Corticostriatal decision making and synaptic plasticity

The spatio-temporal spiking patterns in the above described cortico-striatal-thalamic-cortico (CSTC) circuit have been simulated by modeling the interconnected cortical and subcortical regions as synaptic networks of neuronal assemblies, each comprising spiking neurons of Hodgkin-Huxley type (as described above). Now we describe the set of mechanisms through which these spatio-temporal firing patterns generate the learning driven behaviour (response choice) and feedback driven synaptic plasticity (and thus category learning). Although these equations describe our proposed mechanisms for corticostriatal computation at a much coarseer scale of length and time compared to the physiological scales of spiking neurons and neuroreceptors, they are nevertheless based on well-known physiological and anatomical principles found across the literature.

In principle, all of the following mechanisms can be entirely modeled using only the physiological phenomena at the neuronal length and time scales. However, in this initial rendition of the model we have implemented reasonably simplified population level abstractions of these well defined physiological mechanisms and anatomical constraints. The model is entirely summarized in Table 2. Following is the detailed description of the model components.

**Table 2.** Model Equations

| General form | Specific form |
| --- | --- |
| **Striosome (patch) activity**<br>$\rho_x^{pat}(t, \Delta t) = PAT\left(\sum_{i \in Ctx} W_x^i(t) \cdot [S^i]_{t-\Delta t}^t\right) - LI\left(\sum_{y \neq x} \rho_y^{mat}\right)$<br>$\rho_x^{pat}(t, \Delta t)$ : striosome spiking activity in striatal block $x$<br>$LI$ : Lateral inhibition by other striatal areas | $\rho_x^{pat}(t_n) = H_x(t_n) \sum_{i \in Ctx} W_x^i(t_n) \cdot [S^i]_{t_{n-1}}^{t_n} \quad \vert x \in \{1, 2\}$<br>$H_x(t_n) = \begin{cases} \Theta(\rho_x^{mat}(t_{n-1}) - \rho_y^{mat}(t_{n-1})) & n > 2 \\ 1 & n \leq 2 \end{cases}$<br>$(x, y) \in \{(1, 2), (2, 1)\}$ |
| **TAN activity**<br>$R^{TAN}(t) = Inhibition\left(\sum_x \rho_x^{pat}(t, \Delta t) \middle\vert \kappa^{TAN}\right)$ | $R^{TAN}(t_n) = \begin{cases} \kappa^{TAN}, & n=1 \\ \min\left(\frac{\kappa^{TAN}}{\lambda^{TAN} \sum_x \rho_x^{pat}(t_{n-1})}, \kappa^{TAN}\right), & n>1 \end{cases}$<br>$\kappa^{TAN} = 100, \lambda^{TAN} = 1$ |
| **Matrisome (matrix) activity**<br>$\rho_x^{mat}(t, \Delta t) = MAT\left(\sum_{i \in Ctx} W_x^i(t) \cdot [S^i]_{t-\Delta t}^t\right) + TAN(R^{TAN}) - LI\left(\sum_{y \in N} \rho_y^{mat}(t, \Delta t)\right) \quad \vert y \neq x$<br>$\rho_y^{mat}(t, \Delta t)$ : matrisome spiking activity in striatal block $y$ | $\rho_x^{mat}(t_n) = H_x(t_n) \left[ \sum_{i \in Ctx} W_x^i(t_n) \cdot [S^i]_{t_{n-1}}^{t_n} + Pois(R^{TAN}(t_n)) \right]$<br>$Pois(\lambda)$ : Poisson random variable of mean $\lambda$ |
| **SNc activity**<br>$R^{DA}(t) = Inhibition\left(\sum_x \rho_x^{pat}(t, \Delta t) \middle\vert \kappa^{DA}\right)$ | $R^{DA}(t_n) = \begin{cases} \kappa^{DA}, & n=1 \\ \min\left(\frac{\kappa^{DA}}{\lambda^{DA} \sum_x \rho_x^{pat}(t_{n-1})}, \kappa^{DA}\right), & n>1 \end{cases}$<br>$\kappa^{DA} = 1, \lambda^{DA} = 0.33$ |
| **Action selection and reward feedback**<br>$\mathcal{A} = action_i \quad \left\vert \rho_i^{mat}(t_r) = \max\{\rho_1^{mat}(t_r), \rho_2^{mat}(t_r) \dots\}$ | $\mathcal{A} = \begin{cases} left \ saccade & \vert \rho_1^{mat}(t_r) > 0 \\ right \ saccade & \vert \rho_2^{mat}(t_r) > 0 \end{cases}$ |
| $\mathcal{F}(\mathcal{A}) = \begin{cases} \text{reward} & \vert \mathcal{A}:\text{correct} \\ \text{punishment/no reward} & \vert \mathcal{A}:\text{incorrect} \end{cases}$ | $\mathcal{F}(\mathcal{A}) = \begin{cases} 1 & \vert \mathcal{A}:\text{correct} \\ 0 & \vert \mathcal{A}:\text{incorrect} \end{cases}$ |
| **DA release at corticostriatal synapses**<br>$DA_{SNc} = \int_{t_0}^{t_f} R^{DA}(t) dt + DA^{fb}$ $DA^{fb} : DA^{fb}(\mathcal{F}(\mathcal{A}))$ | $DA_{SNc} = \sum_n [R^{DA}(t_n)] + DA^{fb}$ $\approx [\bar{DA} + [R^{DA}(t_2) - \kappa^{DA}]] + DA^{fb}$ $DA^{fb} = \begin{cases} \theta^{rew} & \vert \mathcal{F}(\mathcal{A}) = 1 \\ 0 & \vert \mathcal{F}(\mathcal{A}) = 0 \end{cases}$<br>**DA<sub>SNc</sub> as a measure of 'Prediction Error'**<br>$DA_{SNc} = \bar{DA} + \epsilon \quad \vert \epsilon : \text{prediction error}$ $\epsilon = \mathcal{R} - \mathcal{Q} \quad \vert \mathcal{R} : \text{reward}, \mathcal{Q} : \text{expectation}$ $\mathcal{R} = [DA]^{fb}, \quad \mathcal{Q} = \kappa^{DA} - R^{DA}(t_2)$ |
| **Corticostriatal synaptic plasticity**<br>$\frac{dW^{ij}}{dt} = F^{ion}(S^i, S^j) \cdot F^{DA}(DA_{SNc}) - \Gamma_{dec}(W^{ij})$<br>*F<sup>ion</sup>*: ionotropic receptor factor, *S<sup>i</sup>*: activity of neuron *i*<br>*F<sup>DA</sup>*: DA receptor factor<br>*Γ<sup>dec</sup>*: synaptic decay | $\Delta W_x^i = H_x \cdot S^i \cdot F^{DA}(DA_{SNc} - \bar{DA}) - \gamma W_x^i \quad \left\vert x \in \{\text{str blocks}\} \right.$ $F^{DA}(y) = K^{DA} \cdot y \cdot (y + \theta_M) \cdot \sigma'(\alpha(y + \theta_M))$ $y = DA_{SNc} - \bar{DA} = \epsilon, \quad \sigma'(x) = \frac{1}{1+e^{-x}} \cdot \frac{1}{1+e^x}$ $K^{DA} = 0.04, \alpha = 2.5, \gamma = 0.01, \theta_M = 1.0$ |
| **Cortico-cortical & thalamo-cortical plasticity**<br>$\frac{dW^{ij}}{dt} = F^{ion}(S^i, S^j) - F^{rev}(DA_{VTA})$<br>*F<sup>rev</sup>*: LTP reversal as function of DA from VTA | $\Delta W^{ij} = K^{ion}[S^i \cdot S^j] \cdot (\bar{W} - W^{ij}) \cdot \mathcal{F}(\mathcal{A})$<br>cortico-cortical : $K^{ion} = 5 \times 10^{-4}$, $\bar{W} = 5$<br>thalamo-cortical : $K^{ion} = 1.7 \times 10^{-5}$, $\bar{W} = 6$ |
| **Thalamo-cortical short-term synaptic augmentation**<br>Synaptic Current<br>$I_{syn}^{ij} = [g_{stp}^i(t) \cdot W^{ij}] \cdot g_{syn}^j(t)(V^i - V_{syn}^j)$ | Conductance Dynamics<br>$\frac{dg_{stp}}{dt} = -\frac{1}{\tau_3} g_{stp} + z \cdot k_{stp}(\bar{G}_{stp} - g_{stp})$ $\frac{dz}{dt} = -\frac{1}{\tau_1} z + g_{\infty}(V)$ $\tau_3 = 2000ms, \tau_1 = 0.1ms, k_{stp} = 0.5, \bar{G}_{stp} = 0.5$ |

### Population activity in STR and SNc

There are four key sub-populations of subcortical regions: *matrisome* (matrix), *striosome* (patch) and *tonically active neurons* (TANs) in the *striatum* and dopaminergic neurons of *substantia nigra pars compacta* (SNc). We consider one sub-population each for TANs and SNc and two competing sub-populations each for matrisome and striosome, respectively part of the two corticostriatal loops described in the circuit model. Since matrix and patch neurons are GABAergic neurons with relatively longer time scales (∼80*ms*), their firings have a cumulative effect over a given time interval. Hence for the striosome and matrisome we consider their cumulative spiking activities *ρ*^*str*^ and *ρ*^*mat*^, respectively. For dopaminergic SNc neurons and cholinergic TANs on the other hand, we consider their instantaneous firing rates *R*^*T AN*^ and *R*^*DA*^ respectively.

#### Striosome activity

Striosome (“patch”) sub-population is driven by spatio-temporal spiking input from cortex, PFC in this case, and is laterally inhibited by matrisomes (“matrix”) of the competing sub-population. The cumulative spiking activity 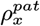 in the striosome sub-population *x* at time *t* over interval Δ*t* is given by

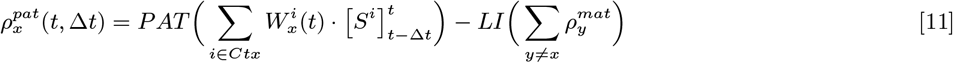

where, *P AT* is a generic function with cortical neuronal activity as input and striosomal (“patch”) activity as output. Here cortical input is represented by 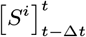 that is the number of spikes occurring in *PFC* neuron *i* during interval [*t* − Δ*t, t*] and 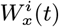 is synaptic weight for the synapse(s) from *PFC* neuron *i* to striatal sub-population *x* (including both patch and matrix). The lateral inhibition function *LI* represents the inhibitory inputs coming from all other areas of the striatum.

For the sake of simplicity, in this paper we used discrete time maps for population activity instead of above described continuous dynamics : *t* ⇒*t*_*n*_ where *t*_*n*_ = *t*_0_ + *n* ·Δ*t*_*inh*_. Here *t*_0_ is the time of stimulus onset and interval Δ*t*_*inh*_ is the time duration of GABAergic inhibition. The discretized form for cumulative striosomal activity (Eq. 11) is given by

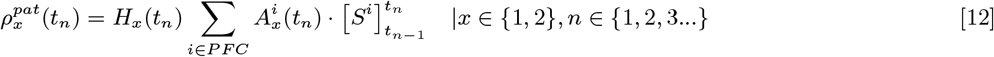

and the simplified form for used for lateral inhibition between two striatal regions modeled here is given by

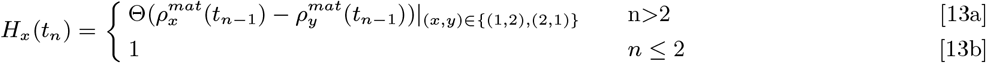

where Θ is the Heaviside function and 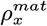 is the cumulative spiking activity of matriosome (“matrix”) subpopulation *x* (see below).

### TAN activity

TANs are driven by intralaminar regions of the thalamus, and are inhibited by all striosomal sub-populations. The firing rate *R*^*T AN*^ can be generally described by:

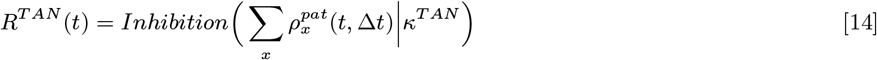

where, *κ*^*TAN*^ is the baseline activity of TAN neurons and *Inhibiton*(*ρ* |*κ*) is a monotonically decreasing function of the cumulative spiking activity *ρ* of the source population, given the baseline spiking activity of the target population is *κ*. The specific discrete form for TAN activity (Eq. 14) we used here is given by

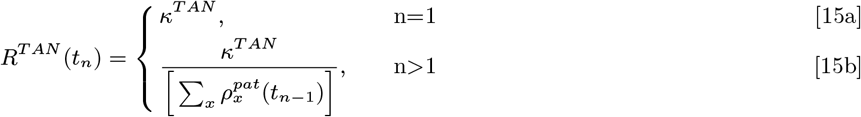

### Matrisome activity

Matrisome sub-populations receive similar inputs as corresponding striosome sub-populations, with an additional modulatory input from TANs. Hence the cumulative spiking activity in the matrisome (matrix) sub-population *x* over interval Δ*t* at time *t* is given by

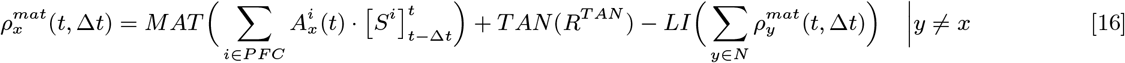

where, *MAT* is a generic function like *PAT* but for the matrisome, with cortical neuronal activity as input and matrisomal (patch) activity as output and *TAN* is the contribution to matrisomal activity from *ACh* modulatory inputs from the TAN neurons. The discrete form of Eq. 16 used in this work is as follows

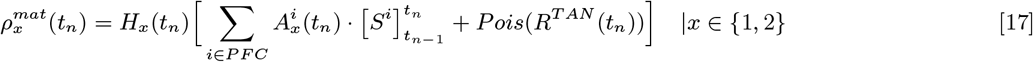

where *Pois*(*λ*) generates a random value from Poisson distribution with mean *λ*.

#### Dopaminergic SNc activity

DA neurons in the SNc, like TANs, are inhibited by GABAergic input from striosomal sub-populations. Their activity can be generally described by

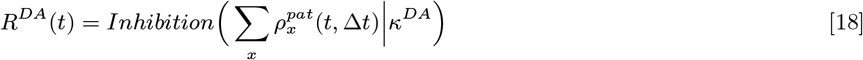

where, *κ*^*DA*^ is the baseline activity of DA neurons in SNc.

The specific discretized form for Eq. 18 used in this work is given by

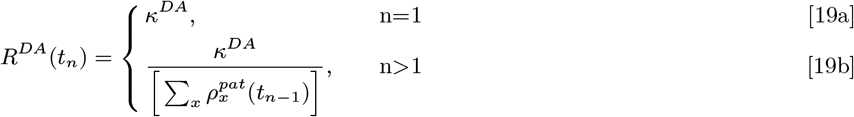

### Action selection and reward feedback

Due to lateral inhibition between striatal sub-populations, one sub-population tends to dominate while others become suppressed. Thus, during a single trial with a given stimulus, only one of the striatal sub-populations consistently remains active, which then activates specific downstream circuits corresponding to a specific response. Thus the response action 𝒜 is chosen at the response time *t*_*r*_ by

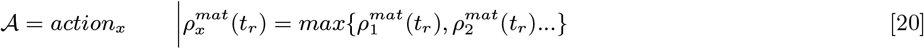

The specific form of selected action is

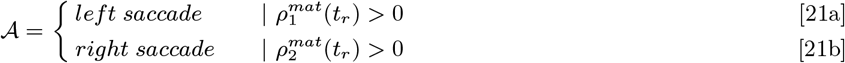

After the response, a simplified feedback ℱ is provided in the form of reward or punishment depending on the correctness of choice

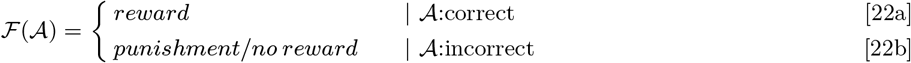

and for this specific case, reward is determined by

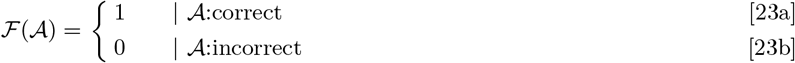

### DA release at corticostriatal synapses

The cumulative amount of dopamine released at the corticostriatal synapses by SNc neurons over the course of a single trial, i.e. from stimulus onset until feedback reward, is given by

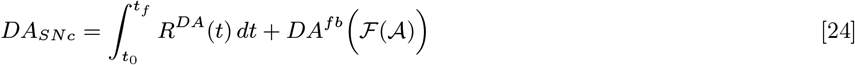

where, *t*_0_ is stimulus onset time, *t*_*f*_(ℱ) is the time of reward feedback and *DA*^*fb*^ F is the amount of DA released due to reward feedback ℱ which is a function of response action 𝒜

The specific discretized form of Eq. 24 used for this work is as follows

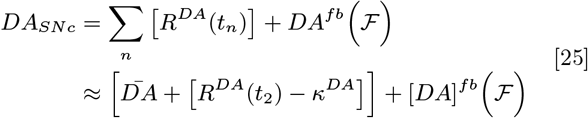

where, 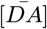 is the baseline DA release in the absence of any striosomal inhibition of SNc.

The strongest suppression of DA neurons from baseline activity as a result of striosomal inhibition occurs at time *t*_2_, since for *t > t*_2_ the lateral inhibition within STR sub-populations suppresses most of the striosomal activity, bringing SNc activity back to baseline. Thus the approximation in Eq. 25 is justified. In this specific case, DA released as a result of feedback is given by:

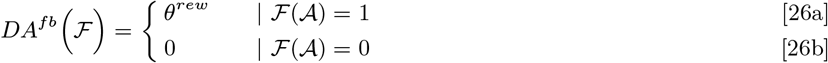

Since the total amount of DA released during a trial directly affects the corticostriatal synaptic plasticity, and thus affects learning, it can also be expressed in a traditional reinforcement learning framework in terms of value prediction error:

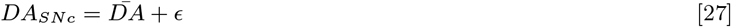

where, error prediction *ϵ* = ℛ − 𝒬 such that the reward is ℛ = *DA*^*fb*^ (ℱ) and the expected reward is 𝒬 = *κ*^*DA*^ − *R*^*DA*^(*t*_2_)

### Corticostriatal synaptic plasticity

Synaptic plasticity on striatal medium spiny neurons (MSNs) follows a set of rules corresponding to the literature on plasticity in basal ganglia (67). A large body of experimental literature indicate three factors involved in the LTP/LTD of corticostriatal synapses (68) : phasic increase in DA release; presynaptic activity; and postsynaptic depolarization. Here in this model we also consider the synaptic decay of corticostriatal synapses. The general form of corticostriatal change is described as :

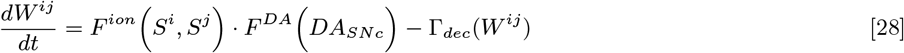

where, *F* ^*ion*^ is the synaptic weight change as a function of pre- and post-synaptic activity of cortical and striatal neurons *S*^*i*^ and *S*^*j*^ respectively, while *F* ^*DA*^ is the D1/D2 receptor factor as a function of the DA from SNc and Γ_*dec*_ is the synaptic decay.

The simplified specific form of Eq. 28 implemented here is given by

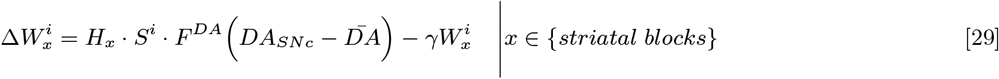

where *H*_*x*_ determines the state of activity in striatal block *x* (see Eq. 13b) and *S*^*i*^ determines the activity in cortical neuron *i*.

The specific form of *F* ^*DA*^ is derived from the experimental literature whose qualitative behaviour is summarized in (68). Due to its qualitative similarity with the BCM curve for synaptic plasticity, even though it explains a completely different set of phenomena, we defined our *F* ^*DA*^ with the same mathematical formas *F* ^*DA*^(*y*) = *K*^*DA*^ ·*y* ·(*y* + *θ*_*M*_) ·*σ*^′^(*α*(*y* + *θ*_*M*_)) where 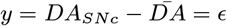 (error prediction). Here the sigmoidal function is defined as 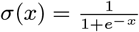.

Due to the specific form of the *BCM* function, it allows both LTP and LTD, depending on the value of the prediction error as quantified by the DA release (see Eq. 27)

### Cortico-cortical and thalamo-cortical synaptic plasticity

In the simulation, specific rules of synaptic potentiation, and reversal of that potentiation, are adopted for synapses in superficial-layer anterior cortical cells, corresponding to consistently replicated studies from the plasticity literature (69–71). Changes in synaptic weights *W* from cortical cell *i* to *j* are described first in general form, illustrating the proposed set of mechanisms to be studied, and then in highly specific form showing a very limited special case of the general form, currently implemented in simulations.

The general form of synaptic change modeled here in is:

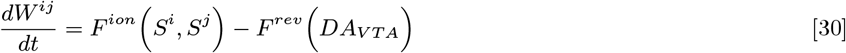

where *F* ^*ion*^ and *F* ^*rev*^ are increments and decrements to synaptic weights; *S*^*i*^ and *S*^*j*^ are the number of spikes from cortical source cells *i* and targets *j*, and *D*_*V TA*_ is the quantity of DA released from the Ventral Tegmental Area (VTA) onto the cortical targets. *F* ^*rev*^ is a function erasing the increment of all recently incremented synapses.

Having identified physiological-level systems mechanisms for potentiation and reversal, the effects of the general form can be substantially simplified for initial incorporation into a simulation. In the currently implemented model, these general forms are implemented as special cases, such that the change to a cortical weight from cell *i* to cell *j* is:

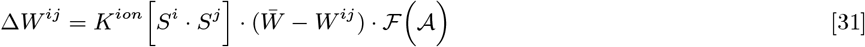

where *K*^*ion*^ is constant and 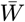 response such that is the maximum value that a weight can attain, while ℱ (𝒜) is a feedback function of the response such that

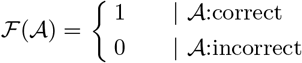

thus capturing the effects of potentiation on correct trials and lack of potentiation on incorrect trials. We apply the same plasticity model for thalamo-cortical synapses as well.

### Short-term augmentation of thalamo-cortical synapses: mechanism for sustained activity

In addition of long term plasticity in the thalamo-cortical synapses explained above, there is an additional short term synaptic augmentation that has been studied in the literature (72, 73). This short-term augmentation of time scales ranging from few milliseconds to few seconds is caused by several factors e.g. the stimulus frequencies and behavioral states of cortical neurons. This short-term synaptic facilitation/augmentation has also been suggested as a mechanism for working memory through persistent reverberation of spiking activity (74). The various molecular mechanisms underlying various types of short-term synaptic augmentation include presynaptic residual calcium (75) and perturbation in postsynaptic firing (76). In our model this short term thalamo-cortical augmentation serves two purposes : (1) to sustain the spiking activity in the circuit even after offset of stimulus and (2) to hold the pre-offset spatial firing pattern in the anterior cortex (“PFC”) intact even during the delay epoch after the offset : the “holding” memory (77). Thus, post stimulus offset, instead of only the spiking activity within the circuit, is sustained but also the “memory” of the stimulus encoded in the spatial firing pattern in the PFC is sustained. This requires a mechanism which augments the thalamo-cortical synapses based on the recent firing of post-synaptic cortical neurons such that the neurons firing during stimulus would be more susceptible to the thalamic excitation even after stimulus offset. We implement this by introducing an extra factor in thalamo-cortical synaptic conductance : 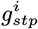, for a synapse from thalamic neuron *j* onto the cortical neuron *i*. The synaptic Equation 3 gets modified for thalamo-cortical connections as:

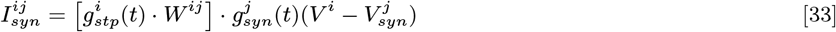

where 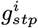, as in the case of synaptic conductance *g*_*syn*_ (see Equations 4), increases due to spiking of neuron *i*:

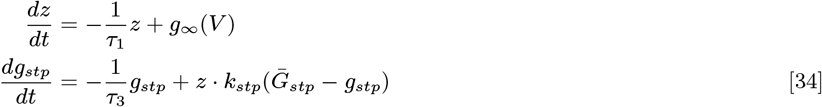

where *τ*_3_ is the decay constant and 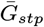 is the asymptotic value (upper bound) of *g*_*stp*_. The intermediate variable *z* is the same in equations 4 and 34.

The equations governing the activity in different kinds of neurons is resummarized in Table 2.

#### Computational Methods

One of the practical difficulties in multi-scale computational neuroscience has been the limitation due to the tension between model flexibility and simulation speed.

Other multi-scale models have been developed based on extensive knowledge of basic physiological and anatomical characteristics of the brain. Existing models can be broadly understood as primarily optimized for cellular-scale mechanisms such as NEURON (78), NEST (79), and Brian (80); or optimized for multi-scale emergent dynamics like The Virtual Brain (81). These platforms present challenges when attempting to produce multi-scale models that span from individual neurons to the whole-brain; they are also not typically constructed to generate extended cognitive operations such as the behavioral perception, learning, working-memory, and decision task presented herein.

To bridge this gap, we developed **Neuroblox**, a computational neuroscience platform optimized for data-driven multi-scale brain modeling (see Methods section in Supplementary Information for further details). Previous modeling efforts to create more realistic and detailed multi-scale biomimetic models have been hindered by the computational intractability of solving massive sets of differential equations. Neuroblox uses the Julia programming language and a variety of computational and mathematical algorithms that lead to orders of magnitude speedups, allowing for hundreds of complex neurons to be simulated in almost real-time. Our computational models are openly available, as stand-alone implementation (github.com/Neuroblox/corticostriatal-circuit-notebook) and also as a model designed in Neuroblox.jl (see https://www.neuroblox.org/). Neuroblox can be used interactively via a graphical user interface.

### Experimental Task Design

The category learning task and the corresponding neurophysiological recording with the macaques (non-human primates, NHPs), on which we specifically model our simulations, is described in detail in (19) and (20) which we summarize here. A single category learning session consists of 600-800 trials where, in each trial, initially the animal has to fixate on a central target onscreen. While fixating, a randomly chosen category exemplar appears on the screen for 600 ms. Each exemplar is a constellation of 7 dots derived from either of the two category prototypes. Each visual stimulus consists of a grid of 15 × 15 pixels lit by seven dots with positions similar to one of two category prototypes (called “A” and “B”) with small random variations. The dots are presented for 0.6 seconds, removed for 1 second (requiring a form of “working memory”), and then a “decision” is made. For the NHPs, the decision was in the form of a saccade to the left (for A) or right (for B) and correct decisions were rewarded with juice. The exemplar epoch (stimulus on) is followed by 1000 ms of delay epoch (stimulus off) in which the animal again fixates on the center and waits for the response cue, after which two saccade targets appear on the left and right of center, each corresponding to one of the categories. The animal responds with a direct saccade to one of the targets indicating the correct category of the exemplar, thus receives a reward (juice drops) on correct response and no reward for incorrect response. The exemplar set for each category is gradually increased starting from 1 for each category, and is doubled in size after each bock of trials. Each block of trials is over once the accuracy of the animal reaches atleast 80% over last 20 trials of that block, after which th next block begins with doubled set of exemplars. Based on this demarcation, first two blocks (100-150 trials) while trials from block 6 onward are considered late learning stage. For every session, a new set of category prototype patterns are used.

### Simulation Protocol

We simulated the category learning trial sessions as performed in the NHPs on our corticostriatal model. The set of random dot pattern stimuli as used in the experiments were also was used in the simulation as the input. Since the visual area in this model does not represent the primary visual area, but some higher area in the hierarchy of visual processing, we added a spatial filter to the dot patterns that approximate the gaussian smoothening as follows. The images are generated in gray-scale with total (15 × 15 =) 225 pixels, where each pixel has brightness value within [0, 1] (0 :black, 1 : white). For each pixel (*i, j*) corresponding to the dot location, the brightness value is 1. For the four adjacent pixels (*i* −1, *j*), (*i* + 1, *j*), (*i, j* −1), (*i, j* + 1), brightness value is 0.6. For all other pixels, brightness is 0. As mentioned in the model description, the pixels map one-to-one on the simulated visual area cortical units, where each unit (neuron) receives a steady input current proportional to the brightness value of corresponding pixel.

In accordance with the experiments, each trial was simulated for 1600 msec, in which the stimulus onset is between 0-600 msec, while during 600-1600 msec there is no visual input. At the end of 1600 msec the circuit model output determines the action choice, and based on the outcome a reward feedback is instantaneously fed into the model (see Model Description above). The synaptic weights are discretely updated after the reward feedback depending on the spatio-temporal spiking patterns within the circuit and the level of dopamine release from SNc and VTA respectively during the trial interval.The next trial cycle begins with the onset of the next visual stimulus. Right from the first trial onwards the stimuli are chosen from either of the two categories randomly out of the entire set, as opposed to gradual doubling of the stimulus set in the experiments (see (20)). For simulations we ran each session for 700 trials, matching the average number of trials in the experiments.

It should be noted that the training regimens are not quite the same: the animal training incorporated i) extensive initial shaping and pilot training, absent from the simulation; and ii) graded introduction of training instances such that initially only single category deformations were shown, and others added subsequently. Since we do not have the trials grouped into blocks based on the size of stimulus samples, we define early and late trials based on their learning behaviour. For simplicity, we kept the group size of early and late trials equal for all simulated sessions viz. 1-150 trials were grouped as early trials and 500-700 trials were grouped as late trials, such that for all the sessions the early trials showed average accuracy below 60% and late trials showed average accuracy above 80%.

### Analysis of Model Physiological Response

#### Field Estimate

We obtain a very simple neural mass activity from the spiking activities of various blocks in the model simulating specific brain regions, viz. Local Spike Summation (LSS), which is simply the spike histogram.

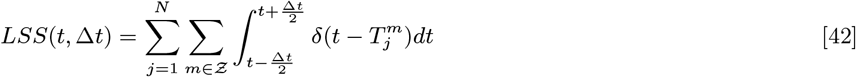

where, 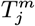is the *m*^*th*^ spike-time for *j*^*th*^ neuron while neuron *j* belongs to a specific block (brain region such as PFC, STR etc.). For all PLV analyses of LSS, we used time bin Δ*t* = 1*ms*. For rastor plot representations in Figures 2 and 3 we used Δ*t* = 20*ms*. We also calculate Average Local Membrane Potential (ALMP) as an alternate surrogate for local field: 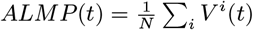 where *N* is the neuronal population size and *V* ^*i*^(*t*) is membrane potential of neuron *i*.

For calculation of Phase Locking Values used to determine synchrony, see the Supplemental Materials.

### Acquisition of incongruent neurons (ICNs)

Incongruent neurons, in both simulations as well as empirical data, were identified during late learning trials as those units (neurons) that were selectively most responsive in the 0 - 200 ms window of stimulus onset during the trials with resultant incorrect outcomes in either of the stimulus categories (A-incorrect and B-incorrect). For testing the predictive capability of incongruent neurons to anticipate incorrect outcomes, we used the first half of late trials for the acquisition of incongruent neurons (acquisition trials) and the second half of late trials for measuring the predictive effect (application trials). Acquisition was done as follows.

1. In order to determine the condition specific preference of single units (neurons) based on their average spiking behavior (as shown in Figure 4A), for each unit (represented by a single row in Figure 4A), we first determine the spiking activity 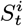 for each unit *i* in response to the respective stimulus of each trial *t*. 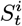 is the number of spikes occurring in unit *i* during a specific time interval (0-200 ms of stimulus onset) for trial *t*. (A notable distinction is the smaller number of average spikes produced in the NHPs compared to the simulation: 1.33 ± 0.05 spikes in 200 msec empirically vs. 9.52± 0.18 spikes per 200 msec in the simulation.)
2. We then calculate normalized spiking activity 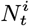 of each unit at a given trial *t* as: 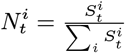, which normalizes the global effects that are unrelated to stimuli but cause trial to trial variation of spiking activity.
3. We sort the acquisition learning trials (first half of late trials) into four conditions based on stimulus category (A vs B) and outcome (correct vs incorrect) viz. A-correct (*C*_1_), A-incorrect (*C*_2_), B-correct (*C*_3_) and B-incorrect (*C*_4_). We then calculate the average normalized activity 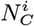 for each unit *i* over group of trials corresponding to each condition *C* where *C* ∈ {*C*_1_, *C*_2_, *C*_3_, *C*_4_}, using 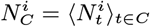.
4. In order to sort the neurons based on their preferred condition, we determine the maximum values of 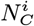 for each unit *I* and we do a second normalization of average normalized activity 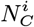 by it’s maximum.

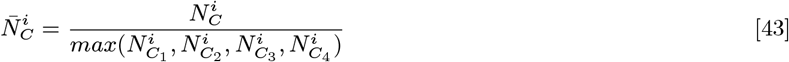

If 𝒩^*C*^ is the set of neurons that prefer condition *C*, we will have 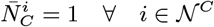.
5. Incongruent neurons (ICNs) are the units that prefer conditions *C*_2_ and *C*_4_ (A-incorrect and B-incorrect), viz. : *ICN* ≡ {𝒩^*C*2^, 𝒩^*C*4^}
6. For empirical data we used an extra clean up step before the identification of incongruent neurons. We had only two criteria : (1) Reject sessions that have insufficient sample size of incorrect trials during the acquisition (≥4 A and B incorrect trials), and (2) Reject the neurons which have too sparse spiking activity throughout acquisition (neurons with 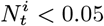 for more than 75% of trials)

### Testing of incongruent neurons (ICNs)

We established the predictive significance of empirical as well as simulated incongruent neurons as follows. After identifying ICNs in the acquisition trials, it was essential to test the statistical significance of predictive accuracy of incorrect choices in the subsequent set of trials, which we called application trials. We hypothesized that the incongruent neurons identified in acquisition trials would continue to be selectively more responsive during A (or B) - incorrect trials during subsequent application trials as well. For a given trial *t*, we use the total normalized activity of condition specific neurons 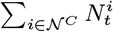 as a decision variable to predict whether the trial is going to be of condition *C*. Thus, given a decision threshold *θ*_*d*_ the trial will be predicted to be A-incorrect if 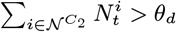, and B-incorrect if 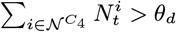. For a fixed *θ*_*d*_, we determine the true positive trials and false positive trials based on the prediction success. If true positive 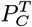 is defined as number of *C* trials accurately predicted to be *C*, true negative 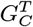 is number of non-*C* trials accurately predicted to be non-*C*, false positive 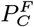 is number of non-*C* trials wrongly predicted to be *C* and false negative 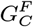 is number of *C* trials predicted to be non-*C*, then true positive rates 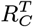 and false positive rates 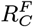 are calculated as : 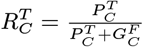 and 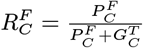. We then used these true positive and false positive rates for conditions *C* ∈ {*C*_4_, *C*_4_} to analyse the predictive significance of incongruent neurons. We performed three separate tests :

#### 1. Group Statistics of ICN prediction

We determined the receiver operating characteristic (ROC) curves (true positive rate vs false positive rate) for each learning session and for each condition type (A-incorrect and B-incorrect) by sweeping across a range of decision threshold values *θ*_*d*_. Predictive performance is estimated using the area under the curve (AUC) which is calculated as area between the ROC curve and the diagonal identity line. It ranges between 0.5 to 0.5. We compared the entire sample of AUC values *A* obtained over all sessions, to the randomized control set of AUCs obtained from using randomly sampled neuron set as predictors of incorrect trials *A*^′^. We calculated the one-sample z-score as 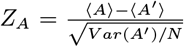, where *N* is the sample size of all ROCs calculated over all sessions. The results are illustrated in Figure 4B.

#### 2. Sample predictive ICN effect

As a specific example to illustrate a simple ICN driven predictive criteria uniformly applicable for all learning sessions, we fix the value of *θ*_*d*_ instead of sweeping it over a range of values as done in the first test described above. For each learning session we determine the value of *θ*_*d*_ for predicting a trial condition *C* (*C* ∈ {*C*_2_, *C*_4_}), from the acquisition trials, using the criteria : 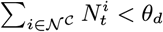 during 75% of the non-*C* trials during acquisition period. Using these acquired values of decision thresholds, we determine the true positive and false positive rates for each condition (A-incorr and B-incorr) across all the sessions. Here predictive performance is measured as difference between true positive and false positive rates : 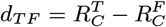. We compare the sample of *d*_*TF*_ obtained from ICNs to the randomized control of predictive performances obtained by randomly sampled neurons used for prediction 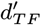. Significance is measured using one sample z-score : 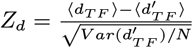, where *N* is the sample size. The results are illustrated in Figure 4C.

#### 3. Predictive real-time statistics of ICN intervention

Based on the predictive criteria suggested in the second test described above, we design a prediction based hypothetical intervention protocol (imagined as a “neuroprosthetic”) during the application trials in real time, so as to test whether there is a significant improvement in the accuracy of the model as well as the animal. The intervention protocol simply is: if the trial is predicted to be incorrect by either A-incorrect 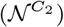 or B-incorrect neurons 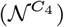, stop the trial/disqualify the response. We calculate the difference of accuracy between intervened behavior and the original behavior Δ. We compare the sample of observed Δ to that obtained from randomized control set of randomly intervened behavior Δ^′^ i.e. randomly selected trials are disqualified instead of trials predicted as incorrect. The significance is measured as one sample z-score: 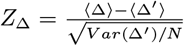, where *N* is the sample size. The results are illustrated in Figure 4D-E

## Acknowledgments

We thank Annabel Driussi for her valuable support with graphics. We thank Anthony Chesebro for useful discussions. The research presented here was funded by the Baszucki Brain Research Fund, United States (LRMP). This work was supported in part by the Office of Naval Research, United States (RG).

## Declarations

Some journals require declarations to be submitted in a standardised format. Please check the Instructions for Authors of the journal to which you are submitting to see if you need to complete this section. If yes, your manuscript must contain the following sections under the heading ‘Declarations’:

## Code availability

All methods shown here, which include an easy-to-use GUI as well as documentation and tutorials, are made freely available through the Neuroblox computational neuroscience platform (neuroblox.org). Neuroblox can be accessed through the JuliaHub repository (juliahub.com).

## Ethics approval

Not applicable.

## Consent to participate

Not applicable.

## Consent for publication

Not applicable.

## Availability of data and materials

Contact the corresponding author.

## Authors’ contributions

Conceptualization: AP, HHS, LRMP, EKM and RG; Data curation: SLB, EGA and EKM; Formal analysis: AP and RG; Funding acquisition: HHS, LRMP, EKM and RG; Investigation: AP and RG; Methodology: AP and RG; Project administration: LRMP; Software: AP, HO, HHS and SS; Supervision: HHS, LRMP, EKM and RG; Validation: AP; Visualization: AP, LRMP and RG; Writing - original draft: AP and RG; Writing - review & editing: AP, SLB, HHS, LRMP, EKM and RG.

## SUPPLEMENTARY MATERIAL

### I. The model has neuron interaction patterns similar to the brain

During the experiment, Simple Visual Cortex Region (SVCR) cells are differentially activated depending on the topographic location of dots in the input stimulus. These neurons will spike (or burst), as a function of voltage summation thresholds in the model (see Methods for corresponding synaptic equations and Figure S1 for an illustration). SVCR cells are only active for 600 ms during presentation of the simulated stimulus (Figure S2A).

**Fig. S1.**
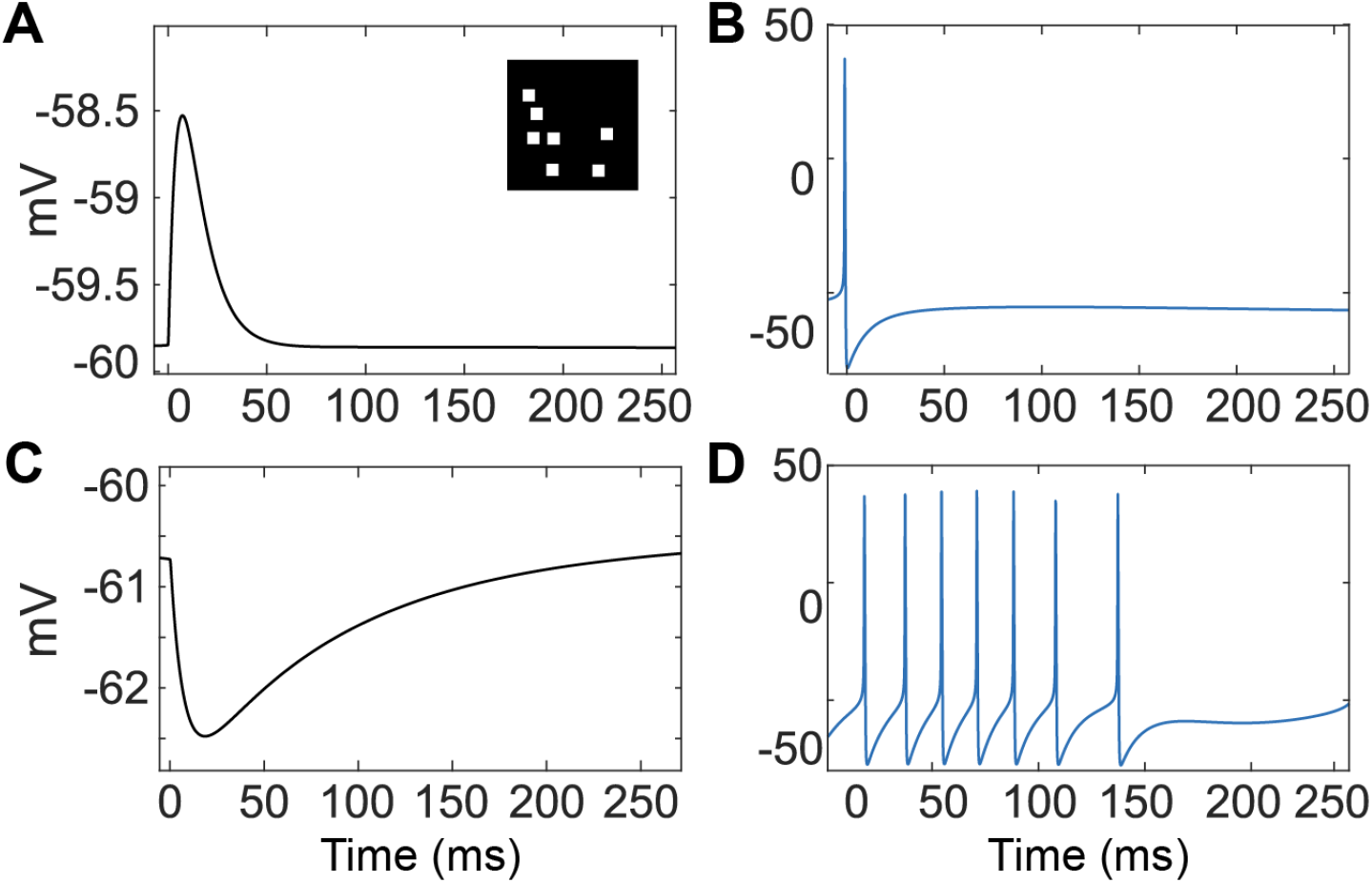
Examples of synaptic potentials calculated for excitatory and inhibitory neurons. (A) Sample Excitatory Post-Synaptic Potential (EPSP); inset: Sample visual stimulus dot pattern. (B) Sample action potential. (C) Sample Inhibitory Post-Synaptic Potential (IPSP). (D) Sample spiking response to visual stimulus

The SVCR inputs are combined with the rhythmic inputs from the simulated ascending (ASC1) brainstem next-gen (NG-NMM) model, so that the predominant response of the SVCR thus exhibits a beta-gamma temporal pattern: brief (40 ms) bursts of high-frequency spiking, separated by approximately 70 ms; i.e., gamma bursts carried on beta rhythms. The activity occurs only on activated input cells in a spatial pattern that is determined by the topographic pattern of the dot-pattern visual inputs. The resulting SVCR activity is input to the simulated anterior cortex (ANTC). These continue to spike due to inputs from ASC1 in a cortico-striatal-thalamo-cortical feedback loop (Figure S2B).

The detailed structure of our model is described in Figure 1 and the Methods. It includes multiple subcircuits whose behavior leads to computational processes. An example of the Striatal Subcircuit and how Tonically Active Neurons (TANs) lead to exploration/explotation is shown in Figure S3.

### II. Calculating and Visualizing Metrics on Multiple Scales

The complex time-variable behavior of our model can be summarized with a few statistical metrics that allow for near-direct comparison to NHP empirical data from (1). Local Spike Summation (LSS) is a simple histogram of the number of spikes across a field of target model neurons (see Methods). The Average Local Membrane Potential is defined as 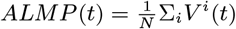, i.e., the average of the membrane potentials (*V* ^*i*^) of each of *N* neurons. While extremely simple, these can be compared to the Local Field Potential (LFP) measured empirically from NHPs (1). These three measures (simulated LSS, simulated ALMP, and empirical LFP) are shown in Figure S4 for both the pre-frontal cortex and the striatum for a single trial. These are represented directly as well as with spectrograms to show the dominant frequencies of activity.

Inspection of Supplimental Figure S4 shows that simulated model spiking and membrane potentials have many similarities to empirical measurements. In the simulated and empirical anterior cortical neurons, spikes are modulated by the rhythmic model ascending input oscillating at a mean rate of 16Hz. At the stimulus offset (600 msec after onset), the response in anterior neurons greatly lessens, but does not go to zero activity, due to cortico-striatal-thalamo-cortical feedback.

In simulated and empirical anterior cortical neurons, there is an initial strong response (150-200 ms) and then a 100 ms silent period. This behavior was first observed in the model and then subsequently confirmed in the empirical data. (We show that the behavior during this first 200 ms also very strongly correlates with final action selection in both the simulation and empirical data.) Further study of this simulation activity is detailed in Supplemental Figure S5

**Fig. S2.**
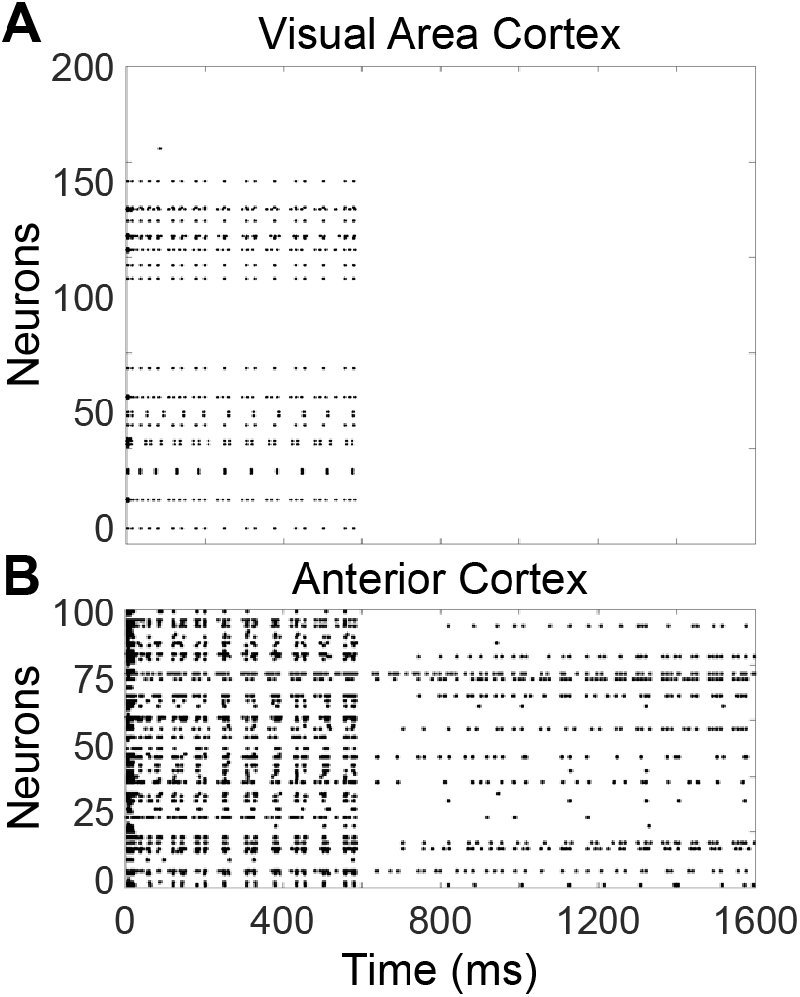
Raster plot showing spiking activity of each simulated excitatory neuron during a single category learning trial within (A) the simple visual cortex region (SVCR) and (B) Anterior Cortex (ANTC) region. The visual stimulus is removed at 600 milliseconds (ms), but a cortico-striatal-thalamo-cortico feedback loop maintains activity in the PFC.

**Fig. S3.**
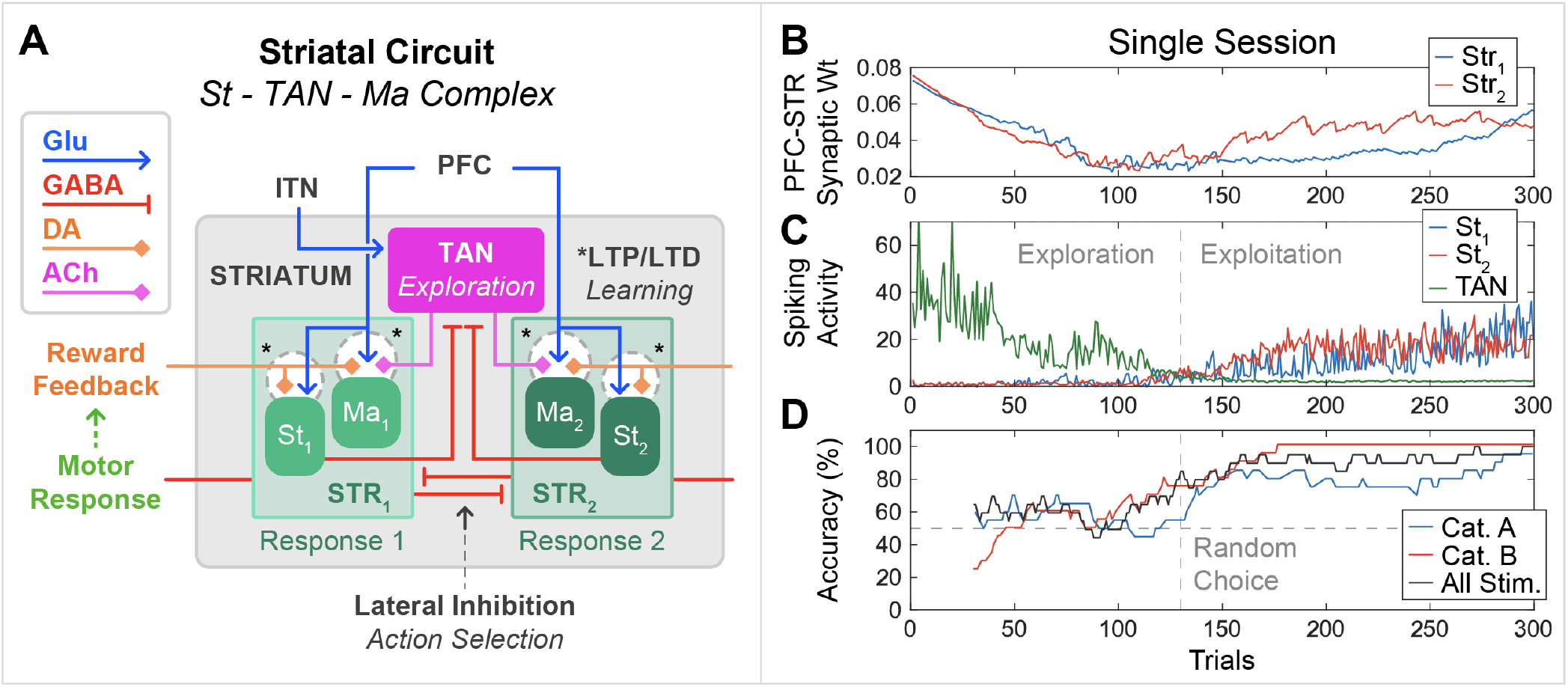
The role of TAN activity in action selection strategy: exploration to exploitation. (A) shows the circuit diagram of striatal complex consisting of matrisome (Ma), striosome (St) and TAN. Cortical inputs are received by Ma and St. St inhibit TAN while TAN modulate Ma. High TAN activity introduces stochasticity in action selection (exploration). Whereas low TAN activity implies action selection is dominated by corticostriatal synaptic weights, thus the strategy is more deterministic (exploitation) (B) Corticostriatal synaptic weight to the two striatal blocks (corresponding to two response choice). They first weaken during early low-performance stage, while strengthens during the late learning stages. The weights do not increase symmetrically for both striatal blocks. (C) Spiking activity (no. of spikes during first 200 ms of stimulus on) in TAN and two STR blocks. When learning stage transitions, TAN activity goes down and STR activities dominate. The transition from exploration to exploitation is thus illustrated. (D) Performance overall (black) and category wise (red and blue.) Performance is also asymmetric in categories, just like synaptic weights in (B)

**Fig. S4.**
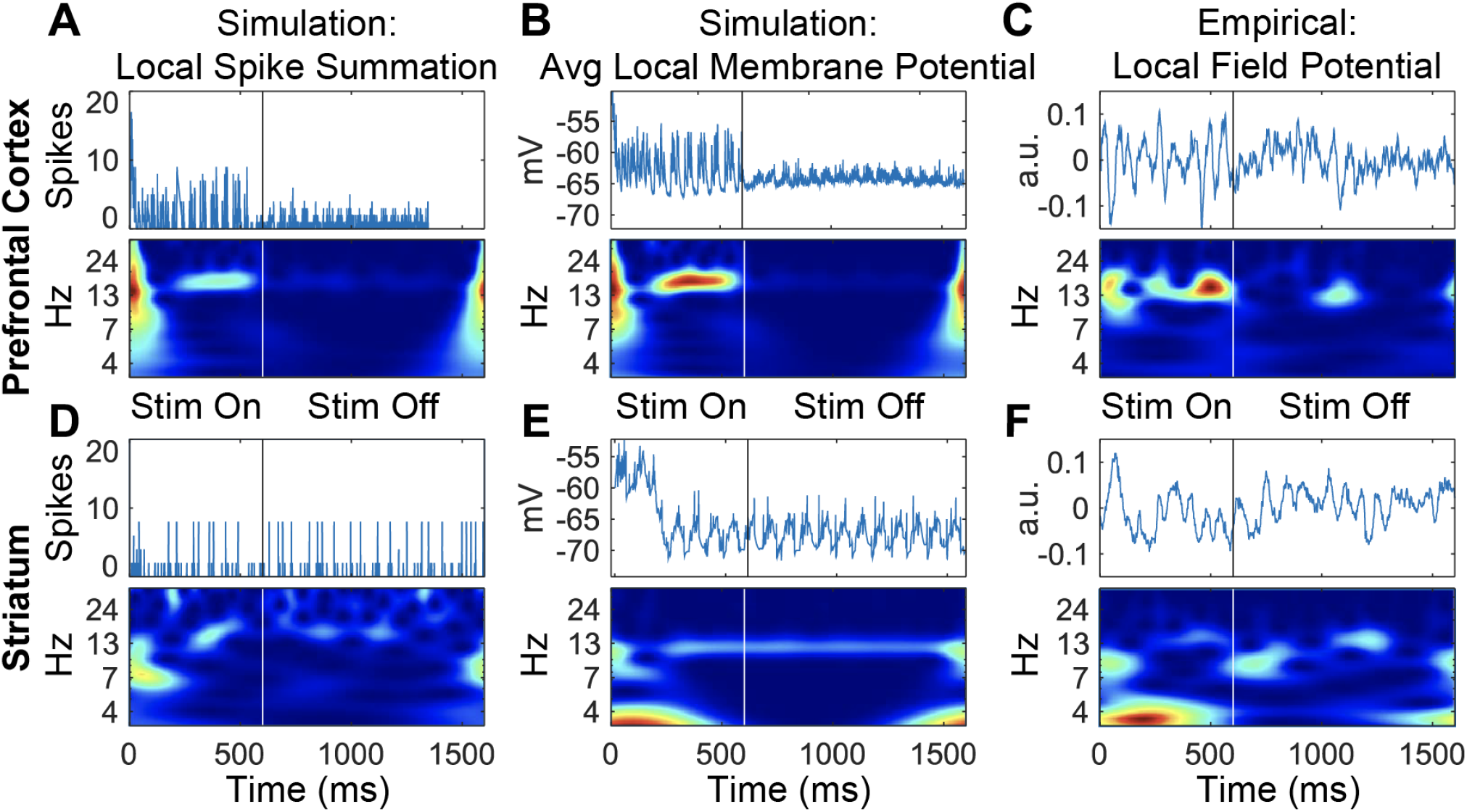
Summaries of neuronal behavior show that model physiologically richly matches empirical data (which was not included in any model training) even for single trials. Each summary shows behavior from the time of stimulus onset (Time = 0) through stimulus offset (600 ms) until final action (1600 ms). The top shows the summary statistic (spikes, millivolts, and arbitrary units) and the bottom shows a spectrogram (see Methods). (A) Local Spike Summation (LSS) histogram of model anterior-cortex neuron spikes. (B) Average local membrane potential (ALMP) straightforwardly calculated by averaging membrane potentials of all cells. (C) Sample empirical local field potential (LFP) measured from macaque cortical electrode. (D) Model LSS response in simulated striatal MSNs. (E) Striatal model ALMP. (F) LFP measured from striatal electrode in macaque. See text for discussion.

A more detailed investigation of this behavior in the model (shown in Figure S5) provided a specific biological mechanism for the post-stimulus silence: the strong initial onset response activates a powerful lateral inhibitory response, building up transient inhibition over the first 50 ms; this in turn strongly inhibits further excitation in the model (for 100 ms) by hyperpolarizing the excitatory neurons (see Figure S5 D and E). The duration of the silent period is thus directly explained by the duration of GABA-Ra receptor-based inhibitory influences, lasting roughly 100 ms. Figure S5 demonstrates that this explains the behavior of the simulation. It is hypothesized that these mechanisms may also directly explain the corresponding post-stimulus-onset silence in the empirical data, which could be confirmed with future investigation. This hypothesis is the first example of how our brain model shows properties similar to the brain that are so specific, that it can provide novel explanations and predictions.

**Fig. S5.**
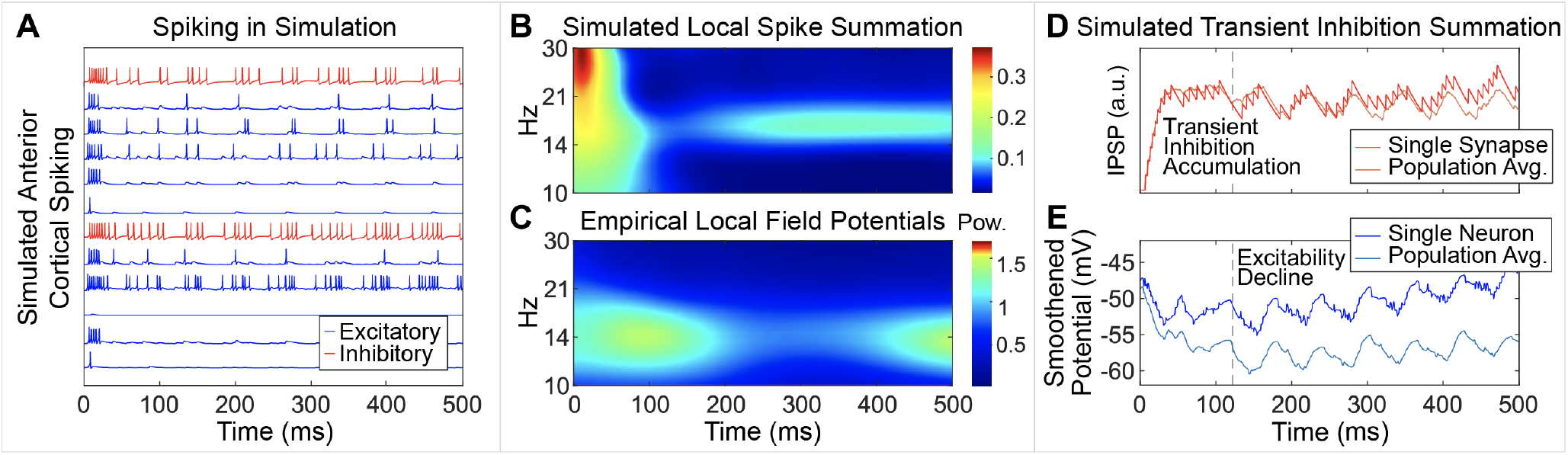
Striatal spiking shows a post-stimulus-onset silence that was subsequently found in empirical data; investigation of the model provides a specific explanation for this silence that makes a novel prediction for future empirical data. (A) Spiking activity in simulated PFC neurons, colored blue for excitatory neurons and red for inhibitory neurons. Note initial burst of spiking activity in a subset of neurons, followed by sparse bursting activity which gradually gets denser. (B) Spectrogram of LSS obtained from simulated spiking activity in the PFC, averaged over all trials in a single learning session, shows transient activity after onset and the following brief silence before further activity. (C) Spectrogram of LFP measured from a sample electrode in anterior cortex (corresponding to PFC), averaged across all trials, shows the same qualitative behavior. (D) Inhibitory synaptic conductivity averaged over entire neural population in simulated PFC shows the accumulation of inhibition during 100 ms after onset. (E) Subthreshold membrane potential averaged over entire population of simulated anterior cortex shows hyperpolarization as a result of the strong transient inhibition, making the neurons less excitable for a brief window, which results in the intermediate period of relative silence seen in (B).

### III. Additional Figures illustrating properties of incongruent neurons

Visual stimuli based on two categories can become increasingly categorized by changing synaptic weights as illustrated in Figure S6. A quantitative calculation of these changes is the “projection angles” between dendritic vectors in the 225-dimensional input space. Learning consists of reweighting that leads to rotation of these vectors, shown in Figure S7. visual stimuli become increasingly categorized int

**Fig. S6.**
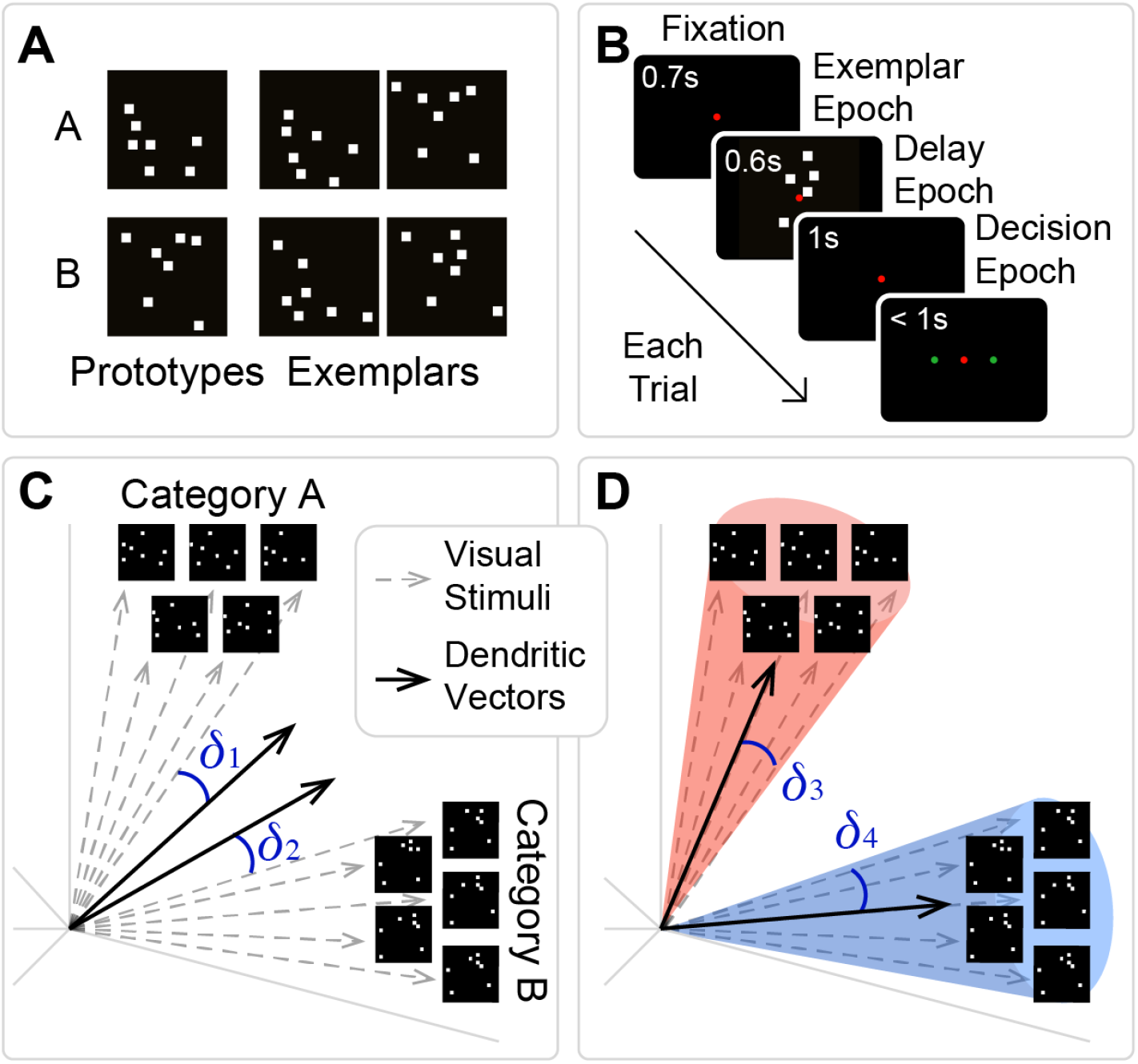
Synaptic increments achieve natural internal clustering of learned image patterns. (A) Two example categories of visual stimuli. Stimuli are the unique exemplars that were created by distortion of the two prototypes. (B) Schematic illustrating the time course of a single learning trial. See references (1, 2) for details of the experimental set-up. (C) Data are random dot patterns that fall naturally into two distinct groups based on their within-group and between-group similarities. They are shown in an input space defined by the abstract coördinates of the possible positionings of dots within each stimulus image. (Although only three coordinate axes are depicted, the input space in fact is 225-dimensional; see Methods.) Because the number of input axons equals the number of potential target synapses, the synaptic dendrite vectors (solid arrows) can be rendered in the same space. The initial positions of synaptic vectors is random with respect to the locations of possible dot-pattern vectors (gray dashed arrows) in the input space. (D) During learning, synaptic weights are modified (via potentiation), causing the dendritic vectors to move toward the mean of the inputs on which the given dendritic vector “wins,” i.e., responds to. After sufficient trials, a given target vector will respond to all and only those input patterns that are closest to the target; this effectively partitions the input space into separate similarity-based categories (red vs. blue shaded areas).

The synaptic plasticity that leads to changing weights also highly aligns individual neurons to specific actions. Figure 4A shows how the spiking activity of individual neurons allows them to be assigned into four categories corresponding to the upcoming choice and outcome: A-correct, A-incorrect, B-correct, and B-incorrect. Figure S8 expands on this categorization by showing how the categorization is related to activity at 200 ms and 1600 ms for the simulated and empirical data in both the cortex and striatum. Figure S9 shows the statistics of spiking frequency averaged over neurons in these categories.

**Fig. S7.**
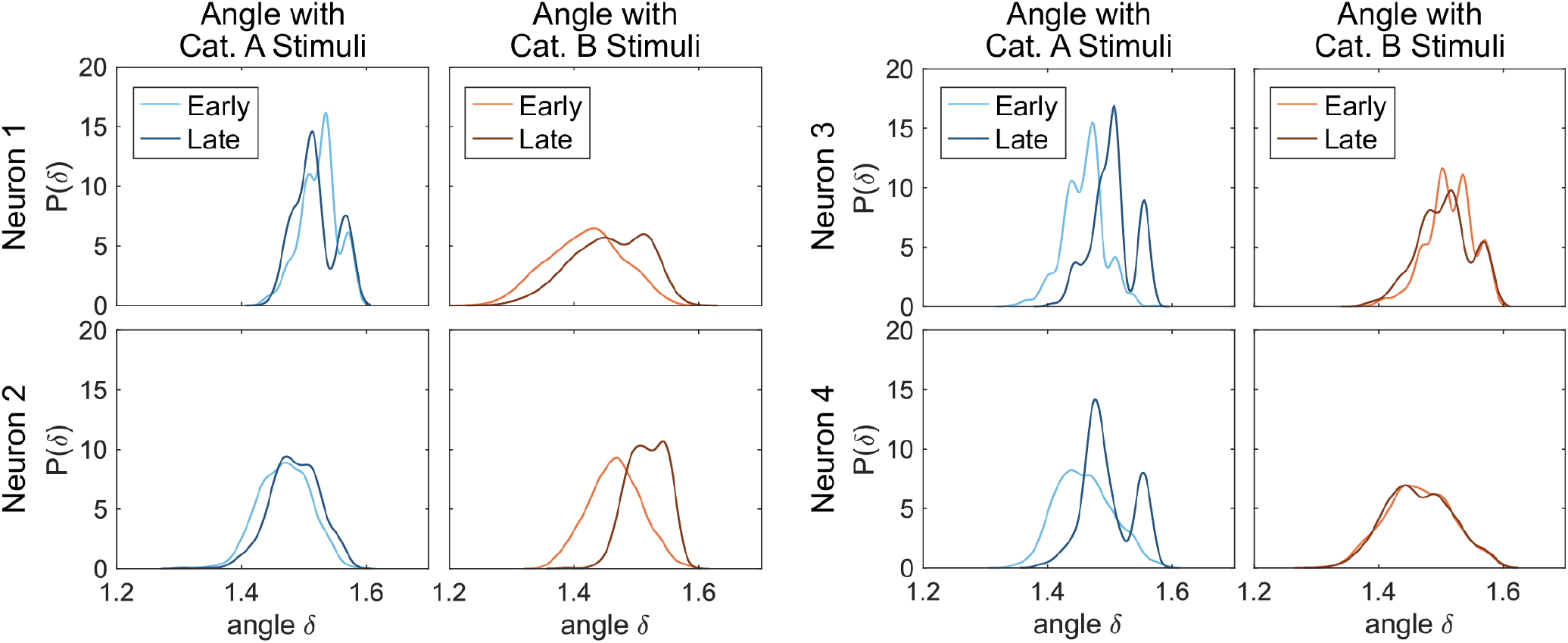
Learning rotates the dendritic vectors of model anterior cortical (PFC) neurons closer to one category and away from the other category. Each panel shows histogram of projection angles between dendritic vector of single neuron and all visual stimulus vectors of a single category (see main text Figure 2). (A) shows two representative neurons whose dendritic vectors rotate towards category A stimuli and away from category B stimuli after the learning session (blue curves in left two panels shift to smaller values while red curves in the right two panels shift to higher values). (B) shows two representative neurons whose dendritic vectors behave the other way i.e. rotate away from category A and towards category B.

**Fig. S8.**
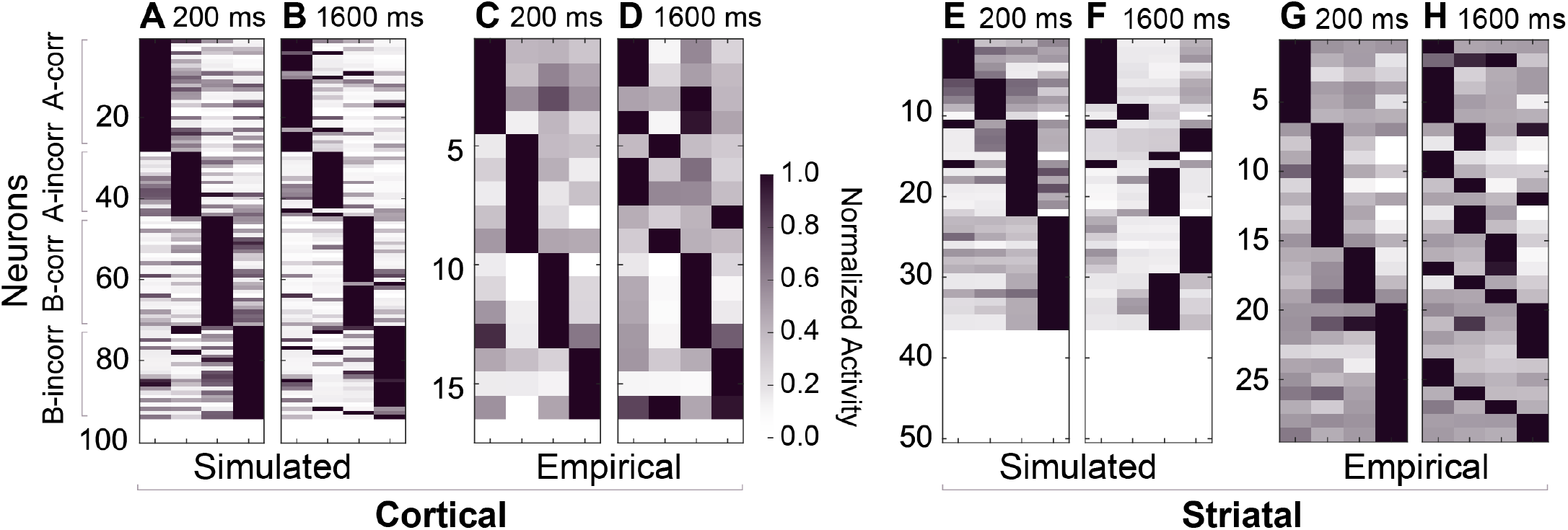
Cortical and striatal cell spiking preferences during late learning trials, modeled and empirical, in four conditions by category and outcome: category A vs. B; outcome correct vs. incorrect. (See vertical axis descriptors at left.) The grayscale shows the activity normalized for that neuron at that time. (A) Simulated cortical spiking preference sorted by four conditions, averaged over first 200 msec. (B) Simulated cortical preference over 1600 msec. (C) Empirical cortical spiking preference in first 200 msec. (D) Empirical cortical preference over 1600 msec. (E) Simulated striatal preference, 200 msec. (F) Simulated striatal preference, 1600 msec. (G) Empirical striatal preference, 200 msec. (H) Empirical striatal preference, 1600 msec. White regions indicate units with no spiking activity.

**Fig. S9.**
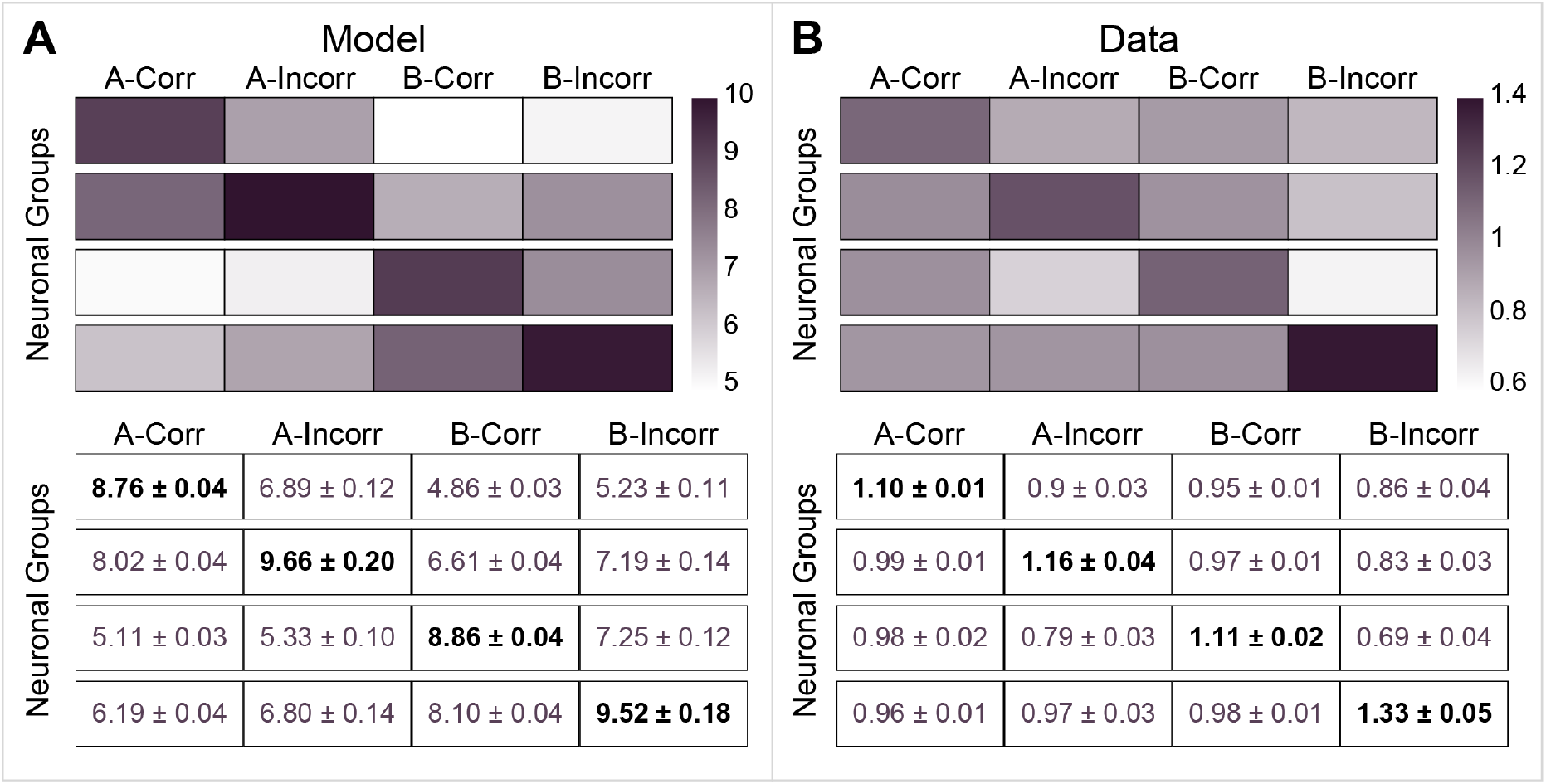
Mean spikes in PFC neurons between 0-200 ms : all sessions. Average spiking activity of four cortical neuron groups as defined by their preferential responsiveness to four types of trials based on category and outcomes for all simulated sessions (A) and for all recorded experimental sessions (B). Row represents a given type of neurons and columns represent their response to a given trial type : A correct, A incorrect, B correct and B incorrect. The tables below the heat maps show the actual numerical values of the average spiking activity and their SEM, in response to the trial during first 200 msec of stimulus onset. Notice the low SEM values indicating the robustness of reliability of the preferential responses.

### IV. Mechanistic explanation for emergence of Incongruent neurons

We then provide a preliminary analysis regarding the emergence of ICNs, viz., even prior to learning, due to the initial synaptic connectivity of ICNs, their firing responses show relatively low discrimination between the two categories as compared to the congruent neurons. Synaptic connectivity of the neurons determines their receptive fields. This can be demonstrated directly in the simulation where precise synaptic connectivity is known and indirectly from the empirically recorded activity from the real neurons in NHPs. In the simulation, a single excitatory anterior cortical neuron can only “see” 18 out of 225 pixels due to the sparse connections to the simulated visual cortex neurons. It is thus possible for category A and B dot patterns (which are random variations around a template) to appear to overlap. (see Figures S6 and S7). Thus the distance between categories, within the subspace of inputs available to a particular neuron, can be quite small, leading to incongruent outputs even though the ensemble of cells together has a high accuracy rate. The limited ability of neurons to “see” is shown in detail in Figure S10 A and B. Examples of the receptive fields of the four categories of simulated neurons are shown while it is also shown quantitatively that “congruent” neurons overlap with more dots and with higher A vs. B selectivity than incongruent neurons. Another indirect estimate of receptive fields of neurons is given by quantitative measure of preference of a neuron for selecting A or B, using the discrimination index defined by (1) and discussed in the Methods. Figure S10 C and D show that “correct” neurons have more biased discrimination indices at the beginning of trials in both the simulated and empirical observations. Thus, neurons with a visual receptive field that starts out biased towards a particular category will evolve to prefer that category highly; incongruent neurons are more likely to start as already ambiguous.

**Fig. S10.**
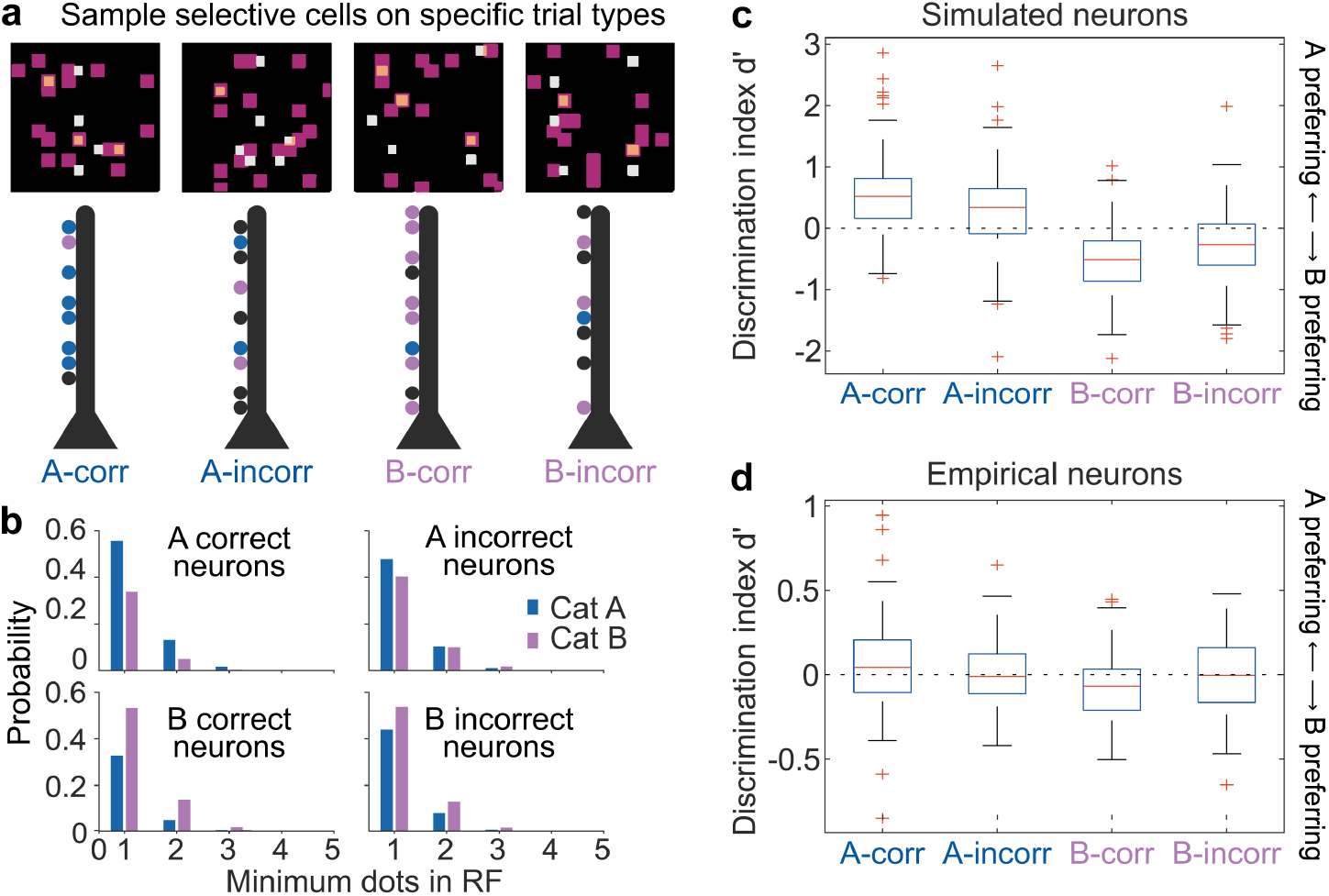
Neuron selectivity for stimuli and for trial outcomes, in simulated and in empirical data. (A) Sample neuron receptive fields (pink shaded regions) for simulated neurons are shown superimposed on sample visual input patterns (white and orange dots : orange dots overlap with the receptive field and white dots lie outside the receptive field), for each of four types (A-correct, A-incorrect, B-correct, B-incorrect). (B) Selective cell responsiveness of simulated neurons are quantified as the probability (given the somewhat random nature of the stimuli) with which stimuli of both categories overlap with the receptive fields (RF) of each of the cell types, where the degree of overlap is given by minimum number of dots occurring within receptive field. (C,D) Calculated discrimination indexes (d’) for the four responsiveness types during early trials, showing d’ values farther from zero for correct A-preferring and B-preferring cells (1st and 3rd columns), but nearer-zero values for responsiveness to A or B on incorrect trials (2nd and 4th columns).

## V. Supplemental Methods

**Table A1.**
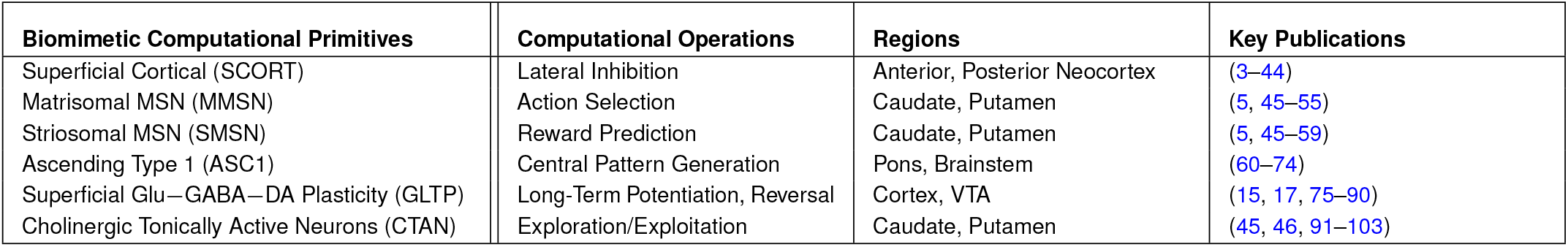
Current Blocks for Corticostriatal Model.

| Biomimetic Computational Primitives | Computational Operations | Regions | Key Publications |
| --- | --- | --- | --- |
| Superficial Cortical (SCORT) | Lateral Inhibition | Anterior, Posterior Neocortex | (3–44) |
| Matrisomal MSN (MMSN) | Action Selection | Caudate, Putamen | (5, 45–55) |
| Striosomal MSN (SMSN) | Reward Prediction | Caudate, Putamen | (5, 45–59) |
| Ascending Type 1 (ASC1) | Central Pattern Generation | Pons, Brainstem | (60–74) |
| Superficial Glu–GABA–DA Plasticity (GLTP) | Long-Term Potentiation, Reversal | Cortex, VTA | (15, 17, 75–90) |
| Cholinergic Tonicly Active Neurons (CTAN) | Exploration/Exploitation | Caudate, Putamen | (45, 46, 91–103) |

### V.A. Detailed model description

#### Extensive anatomical and physiological features incorporated into the present simulation

A sizeable body of references was used in the conception and construction of the several subcircuits and neural assemblies that constitute the present brain circuit simulation (Table A1). These consist of attributes that are highly cited in the literature.

#### Neuronal Dynamics

Each neuron is modeled using ionic conductance based point neuron models, with the Hodgkin-Huxley formalism (104). An overview of this behavior is shown in the Methods with specific details discussed here. For simplicity, in this model we consider only voltage gated Sodium and Potassium channels. The membrane potential is described by:

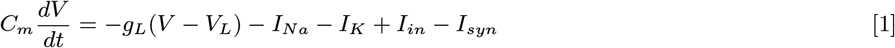

where membrane capacitance *C*_*m*_ = 1*µ* F *cm*^−2^, leak conductance *g*_*L*_ = 0.1 mS *cm*^−2^, *I*_*in*_ is the input current, which can either be external stimulus or background activity, and *I*_*syn*_ is the synaptic input. Sodium current 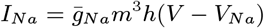 and potassium current 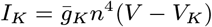 are driven by gating variables {*m, h, n*} whose dynamics are governed by the general form :

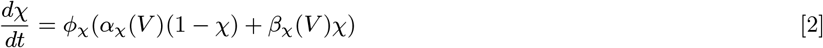

Here, *χ* is the gating variable and *ϕ*_*χ*_ is the temperature related effect on the time scale.

The following parameters were used as shown in ref. (105) : 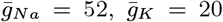, *V*_*Na*_ = +55*mV, V*_*K*_ = −90*mV, V*_*L*_ = −60*mV* and *ϕ* = 5.

Different threshold values for used for pyramidal and inhibitory cells. For excitatory pyramidal cells we used the following (105):

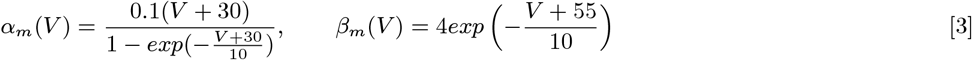

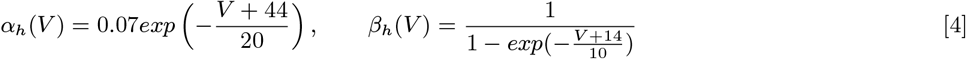

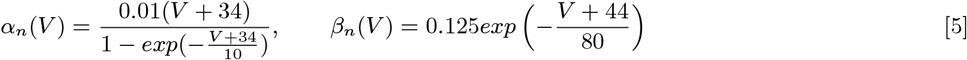

Similarly, for inhibitory neurons we used the threshold values implemented in ref. (106):

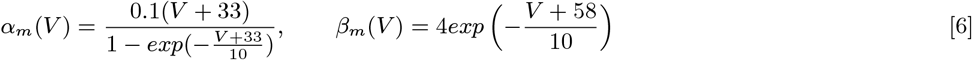

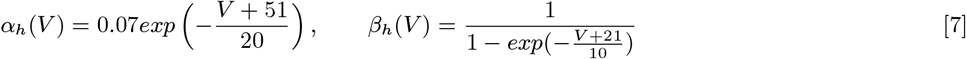

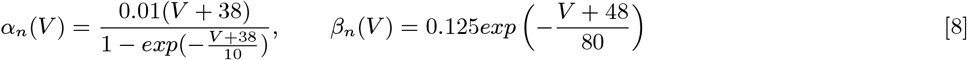

### Synapse

Synaptic current in eq. 1 for a neuron *i* is the sum of synaptic inputs from all incoming axons {*j*} viz. 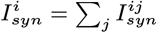. Synaptic current from neuron *j* to neuron *i* is given by

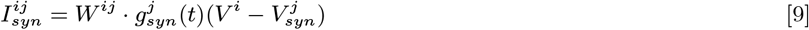

where *W* ^*ij*^ is the synaptic weight from neuron *j* to *i*, a quantity which estimates density of dendritic spines and is modified by Long Term Potentiation (LTP) and Long Term Depression (LTD). 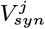 is synaptic reversal potential for presynaptic neuron, which for excitatory neuron is 0*mV* and for inhibitory neuron is −70*mV*. The synaptic conductance 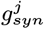 is a function of spiking activity of the presynaptic neuron. As suggested in reference (107), synaptic conductance *g*_*syn*_ is one of the state variables for each neuron whose dynamics is described by damped oscillator equation:

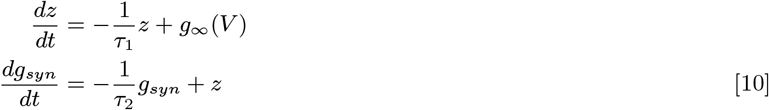

which is driven by impulse force *g*_∞_(*V*) defined as follows (108)

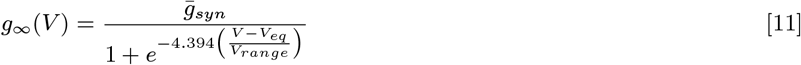

For excitatory neurons 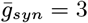, *V*_*eq*_ = 10*mV* and *V*_*range*_ = 35*mV* and for inhibitory neurons 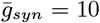, *V*_*eq*_ = 0*mV* and *V*_*range*_ = 35*mV*.

Equations 10 provide two different time scales *τ*_1_ and *τ*_2_ for the synaptic conductance waveform which determines the rate of rise and decay respectively. For excitatory neuron we set *τ*_1_ = 0.1 ms and *τ*_2_ = 5 ms while for inhibitory neurons, we used *τ*_1_ = 0.1 ms and *τ*_2_ = 70 ms.

#### Ascending Systems and Intralaminar Thalamic Nuclei

The simulated ascending system (ASC1) is an extreme simplification of ascending systems such as the cholinergic basal forebrain and noradrenergic locus coeruleus; it produces rhythmic activity via a neural mass model. That activity becomes simulated modulatory input to cortex, driving a beta rhythm in the cortex (63). The ASC1 simulation thus functions as a central pattern generator. It has been modelled using a simple ‘Next Generation’ neural mass model (109, 110) which is a mean field approximation of a network of rhythmic neurons (111). A single neural mass representing an ascending system nucleus comprises a synaptically coupled population of excitatory (E) and inhibitory (I) neurons. Their collective activity is defined by the complex valued Kuramoto order parameter *Z*_*e*_ and *Z*_*i*_ respectively, while their synaptic interaction is measured by synaptic conductivities *g*_*ee*_, *g*_*ei*_, *g*_*ie*_ and *g*_*ii*_ such that *g*_*ab*_ is the synaptic conductivity from population *b* to population *a*, where *a, b* ∈ {*E, I*}. The dynamics is described as :

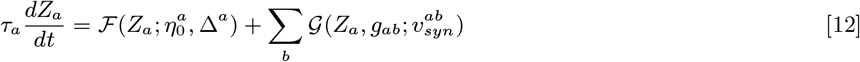

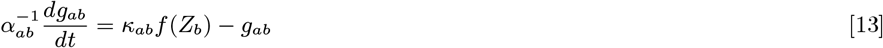

where

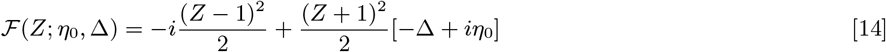

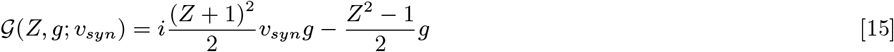

and the population firing rate *f* is given by

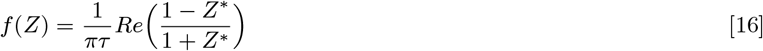

The firing rate *f* (*Z*_*e*_) of population *E* directly affects the modulation of the target neurons, in the form of an input current to the target neurons such that *I*_*mod*_ = *k*_*LC*_*f* (*Z*_*e*_). In this simulation, those targets are simulated somatostatin feed-forward inhibitory cortical neurons.

For ASC1, we used the following parameters : *τ*_*e*_ = 52, *τ*_*i*_ = 26,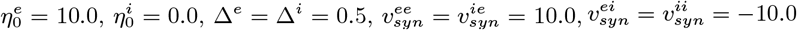, *α*_*ee*_ = *α*_*ie*_ = 0.385, *α*_*ei*_ = *α*_*ii*_ = 0.031, *κ*_*ee*_ = *κ*_*ii*_ = 0, *κ*_*ei*_ = *κ*_*ie*_ = 15.6, *k*_*LC*_ = 44.

Similarly, Intralaminar Thalamic Nuclei (ITN) were modeled as a neural mass which targets the TAN neurons (112) which then drives the alpha rhythm (12 Hz) in the striatum. We again use the next generation neural mass model with the following parameters : *τ*_*e*_ = 72, *τ*_*i*_ = 36, 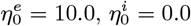, Δ^*e*^ = Δ^*i*^ = 0.5, 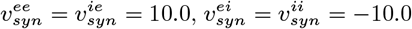, *α*_*ee*_ = *α*_*ie*_ = 0.278, *α*_*ei*_ = *α*_*ii*_ = 0.022, *κ*_*ee*_ = *κ*_*ii*_ = 0, *κ*_*ei*_ = *κ*_*ie*_ = 21.6, *k*_*ITN*_ = 100.

Ascending systems are known to substantially participate in cortical rhythmicity, via modulatory inputs (113) either to pyramidal cells or interneurons (60–67). The resulting rhythmicity, in multiple EEG frequency bands, shows evidence of both intrinsic and extrinsic influences (114–117). Differential effects of intrinsic and ascending components may be studied in part by modeling them separately; we here specifically investigate the extent to which simple rhythmic driving from ascending signals can give rise to complex effects of behavioral states quite beyond those of simple rhythmicity: for instance, the phase, amplitude, and synchrony of these rhythms and their dependencies on different behavioral states.

In Parkinson’s disease, evidence suggests that symptoms are accompanied by changes to the striatal beta rhythm with possible involvement of the subthalamic nucleus and globus pallidus pars interna (GPi) (118–120). The potential applicability of the present work to Parkinson’s and other clinical conditions is of great interest.

The extraordinary anatomical range and physiological driving power of modulatory ascending systems targeting cortical and subcortical structures suggests their involvement in rhythmic activity; their implication in clinical disruptions is a subject of ongoing study.

We model a simplified ascending system as a next-gen neural mass model (NG-NMM) (109) with parameters set such that calculated local membrane potentials exhibit increased power in the driving frequency range as a Gaussian centered at 16 Hz. The cholinergic basal forebrain, as one ascending example, provides two cell populations with two sets of ascending fibers: cholinergic axons contacting pyramidal cortical cells plus inhibitory axons that contact somatostatin inhibitory feedforward interneurons in cortical and subcortical targets (61, 62). By contrast, the locus coeruleus (LC) reportedly projects noradrenergic axons to pyramidal cells; however, there also exists a locus coeruleus-adjacent population of inhibitory cells that reciprocally contact LC neurons, and these inhibitory cells ascend to possible somatostatin targets in cortex (121). Although these LC-adjacent cells are not typically considered a part of LC proper, it is possible that this organization may reflect something analogous to that in basal forebrain. Although brainstem nuclei are typically excluded from human brain imaging, recent work is in accord with extensive prior evidence, indicating extensive alignment of cortical and brainstem activity (122). In the present simulations, an abstracted ascending system is implemented as a stand-in, with the intention of initially exploring the simplest potential effects of such systems.

The basic rhythmic input is thus modeled at present by the next-generation neural mass models (NG-NMM). Notably, all other rhythmic effects in the present work: corticostriatal synchrony; phase-locking values (PLV); learning-related frequency and synchrony changes; rhythmic changes accompanying behavioral states (stimulus, working memory delay, action selection, etc.), all arise solely from modeled unit neuronal activity and synaptic change in the simulation. It is hypothesized that these effects occur as identified in the present simulation, and further elaboration of rhythms will not fundamentally alter these primary findings.

### Corticostriatal decision making and synaptic plasticity

The spatio-temporal spiking patterns in the above described cortico-striatal-thalamic-cortico (CSTC) circuit have been simulated by modelling the interconnected cortical and subcortical regions as synaptic networks of neuronal assemblies, each comprising spiking neurons of Hodgkin-Huxley type (as described above). Now we describe the set of mechanisms through which these spatio-temporal firing patterns generate the learning driven behaviour (response choice) and feedback driven synaptic plasticity (and thus category learning). Although these equations describe our proposed mechanisms for corticostriatal computation at a much coarseer scale of length and time compared to the physiological scales of spiking neurons and neuroreceptors, they are nevertheless based on well known physiological and anatomical principles found across the literature.

A notable feature of corticostriatal projections is the distinction between cortical intratelencephalic (IT) and pyramidal tract (PT) neurons; the former project ipsi- and contralaterally to the striatum whereas the latter project only ipsilaterally to the striatum and other structures (123). Moreover, IT and PT neurons exhibit distinct local cortical circuit arrangements and distinct physiological characteristics. Possible differential effects of distinct dopamine receptors has been reported but remains uncertain (124–126).

In principle, all of the following mechanisms can be entirely modelled using only the physiological phenomena at the neuronal length and time scales. However, in this initial rendition of the model we have implemented reasonably simplified population level abstractions of these well defined physiological mechanisms and anatomical constraints. The model is entirely summarized in Table 2, Main Text. Following is the detailed description the model components.

This general form partially captures a set of physiological phenomena from the literature, directly relating to experimentally validated synaptic change:

- Brief bursts of high-gamma (∼80-100 Hz) spiking, recurring at intervals of 5 Hz (thus, “theta-burst” or “theta-gamma” patterns) induce very long-lasting synaptic potentiation; this phenomenon has been demonstrated to various extents in hippocampal field CA1, and in posterior and anterior neocortical regions. The mechanisms for this are surprisingly well worked out and extensively replicated (see, e.g., (76–81)).
- Reciprocally, individual spikes that recur at 5 Hz intervals (“theta-spike” patterns) cause very recently-induced LTP to be reversed; this adenosine-dependent phenomenon is linked to interference with consolidation of recent LTP, which otherwise remains fragile for several minutes after induction (75, 127).
- In a cortico-striatal-thalamic-cortical simulation, ongoing bursting, at 14-16 Hz repetitions (beta-burst), is induced by correct behavioral responses to learned stimuli. As described, this low-beta-burst stimulation gives rise to synaptic potentiation in cortex, strengthening responses to learned cues.
- During early training, when an incorrect behavioral response to a given stimulus was made, the mismatch triggered dopamine responses, both in the SNc and in the VTA. The SNc response participates in feedback to striatal MSNs, as part of a reinforcement step that causes changes to corticostriatal synapses. This SNc dopamine response is in direct concordance with the widely-agreed upon and highly cited findings of many studies on reinforcement learning in the striatal complex (56–59, 128)..
- Meanwhile, the VTA dopamine response is far less well defined in the literature than the SNc response; the simulation implements a well-cited effect of dopamine in prefrontal cortex, in which dopamine activity suppresses cortical excitatory cell gamma bursting and instead allows cortical single spikes (e.g., (82, 83)), i.e., in this case, spiking recurring at beta.
- Thus, incorrect behavioral responses generate beta-spiking, rather than beta-bursting. These incorrect behaviors thereby cause reversal of the potentiation that otherwise would have taken hold in the cortical responses (75, 127), thus preventing the incorrect encoding from being permanently learned. It is notable that this confluence of ascending systems together with rhythmic bursts and spikes, yields a result that can (surprisingly) be interpreted at the level of specific receptor activity on individual synapses.

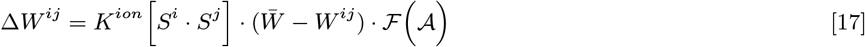

where *K*^*ion*^ is constant and 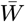 response such that is the maximum value that a weight can attain, while ℱ (𝒜) is a feedback function of the

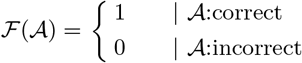

#### V.B. Computational Methods

One of the practical difficulties in multi-scale computational neuroscience has been the limitation due to the tension between model flexibility and simulation speed. For this reason, we chose to code our mathematical models in Julia, which shows orders of magnitude speed improvements in ODE systems compared to Python and MATLAB(129).

Our neural circuits are modeled using the Julia ModelingToolkit (MTK) component-based block-modeling system (130). MTK generates code using a symbolic preprocessing system that allows for automating transformations such as index-reduction of differential-algebraic equations via Pantelides algorithm (131, 132), alias elimination, tearing of nonlinear systems, symbolic simplification (133, 134), and more to produce differential equation and nonlinear systems which have improved numerical stability over naive approaches. In addition, MTK uses sophisticated algorithms such as the D* algorithm (135) to generate system Jacobians in asymptotically less time than traditional techniques based on symbolic, automatic (136), and numerical differentiation. Code generation improvements are coupled with advancements in the numerical solving of differential equations provided by the Julia-based DifferentialEquations.jl methods (137). In particular, we created a MTK library of modular computational dynamical building blocks that consists of hierarchical composites of indiviual Hodgkin-Huxley neurons (cortical blocks), next-generation neural mass models, and various reinforcement computational primitives that can be assembled using a directed graph structure. Once all the connections are defined, we transform the directed graph into a set of symbolic Ordinary Differential Equations (several thousand ODEs) that are then symbolically simplified and compiled into numerically optimized functions to interface with the Julia DifferentialEquations.jl ODE solver packages. We chose to use the Vern7 solver (Verner’s “Most efficient” 7/6 Runge-Kutta method (138)) to optimize both speed and accuracy of the solution. For our model system with 600 Hodgkin-Huxley neurons, simulation of spiking activity over a single trial duration of 1600 ms, takes only 50-60 seconds.

Our computational models are openly available, as stand-alone implementation (github.com/Neuroblox/corticostriatal-circuit-notebook) and also as a model designed in Neuroblox.jl (Neuroblox.org). (Within this system, ModelingToolkit.jl automatically transforms equations into simpler forms for numerical solving, and the system can be used interactively via a graphical user interface.)

#### V.C. Calculating Phase Locking Synchronization (PLV)

Wavelet analysis of the Local Spike Summation (LSS) and subsequent calculation of synchrony measure Phase-Locking Value (PLV) was performed using the same methods as was followed in the measured LFPs in reference (1). First, the wavelet transform of each LSS timeseries was computed by convolution of LSS with Morlet wavelet (139) at 5 octaves between 2-64 Hz at frequency resolution of 0.1 octave. For this analysis, we used MATLAB based software available at http://atoc.colorado.edu/research/wavelets/. The Wavelet transform of a single LSS gives a phase time series *ψ*(*t*) for every frequency *f*. Frequency specific *PLV* between two LSS timeseries is given by 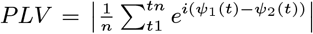. Two perfectly phase-synchronised timeseries would measure a PLV as 1 while completely asynchronous time series would give PLV as 0. As done in the neurophysiological analysis in (1), we calculate the PLV over the last 500 ms of exemplar (stimulus on) and delay (stimulus off) epochs, viz. between 100 to 600 ms and between 1100 to 1600 ms respectively. In order to compare the PLV values of simulated LSS with that of the empirical LFPs in reference (1), we subtracted from the observed PLVs the mean PLV obtained from randomised surrogate ensemble. The surrogate ensemble was obtained by randomising the trial orders of the LSS time series from PFC ans STR and we took 1000 realisations of these randomisations.

Differential synchrony behaviour between two sets of trials (correct and incorrect) is measured by the discrimination index *d*^′^ as also used in (1). The discrimination index is the difference between mean PLVs of two sets of trials normalised by their combined standard deviation : 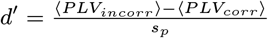, where 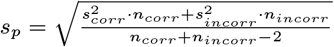. We also remove he sampling bias by converting the observed discrimination *d*^′^ into Z-scores with respect to another randomised surrogate ensemble. This ensemble is obtained by randomly shuffling the trials between two sets and then calculating *d*^′^. We used 400 realisations of randomised surrogates for *d*^′^.

## References

1. GMG Shepherd, Corticostriatal connectivity and its role in disease. Nat. Rev. Neurosci. 14, 278–291 (2013).

2. M Bekhbat, et al., Functional connectivity in reward circuitry and symptoms of anhedonia as therapeutic targets in depression with high inflammation: evidence from a dopamine challenge study. Mol. Psychiatry 27, 4113–4121 (2022).

3. BI Rappaport, S Kandala, JL Luby, DM Barch, Brain Reward System Dysfunction in Adolescence: Current, Cumulative, and Developmental Periods of Depression. The Am. J. Psychiatry 177, 754–763 (2020).

4. E Roltsch Hellard, et al., Optogenetic control of alcohol-seeking behavior via the dorsomedial striatal circuit. Neuropharmacology 155, 89–97 (2019).

5. S Bariselli, DM Lovinger, Corticostriatal Circuit Models of Cognitive Impairments Induced by Fetal Exposure to Alcohol. Biol. Psychiatry 90, 516–528 (2021).

6. K Sabaroedin, J Tiego, A Fornito, Circuit-Based Approaches to Understanding Corticostriatothalamic Dysfunction Across the Psychosis Continuum. Biol. Psychiatry 93, 113–124 (2023).

7. EM Izhikevich, GM Edelman, Large-scale model of mammalian thalamocortical systems. Proc. Natl. Acad. Sci. 105, 3593–3598 (2008).

8. V Jirsa, et al., Personalised virtual brain models in epilepsy. The Lancet Neurol. 22, 443–454 (2023).

9. A Leblois, T Boraud, W Meissner, H Bergman, D Hansel, Competition between Feedback Loops Underlies Normal and Pathological Dynamics in the Basal Ganglia. J. Neurosci. 26, 3567–3583 (2006).

10. K Gurney, TJ Prescott, P Redgrave, A computational model of action selection in the basal ganglia. I. A new functional anatomy. Biol. Cybern. 84, 401–410 (2001).

11. K Gurney, TJ Prescott, P Redgrave, A computational model of action selection in the basal ganglia. II. Analysis and simulation of behaviour. Biol. Cybern. 84, 411–423 (2001).

12. MJ Frank, Dynamic dopamine modulation in the basal ganglia: a neurocomputational account of cognitive deficits in medicated and nonmedicated Parkinsonism. J. Cogn. Neurosci. 17, 51–72 (2005).

13. MJ Frank, Hold your horses: a dynamic computational role for the subthalamic nucleus in decision making. Neural Networks: The Off. J. Int. Neural Netw. Soc. 19, 1120–1136 (2006).

14. JJ DiCarlo, DD Cox, Untangling invariant object recognition. Trends Cogn. Sci. 11, 333–341 (2007).

15. JJ DiCarlo, D Zoccolan, NC Rust, How does the brain solve visual object recognition? Neuron 73, 415–434 (2012).

16. MD Humphries, RD Stewart, KN Gurney, A Physiologically Plausible Model of action Selection and Oscillatory Activity in the Basal Ganglia. J. Neurosci. 26, 12921–12942 (2006).

17. FG Ashby, JM Ennis, BJ Spiering, A neurobiological theory of automaticity in perceptual categorization. Psychol. Rev. 114, 632–656 (2007).

18. FG Ashby, MJ Crossley, A Computational Model of How Cholinergic Interneurons Protect Striatal-dependent Learning. J. Cogn. Neurosci. 23, 1549–1566 (2011).

19. EG Antzoulatos, EK Miller, Differences between Neural Activity in Prefrontal Cortex and Striatum during Learning of Novel Abstract Categories. Neuron 71, 243–249 (2011).

20. EG Antzoulatos, EK Miller, Increases in Functional Connectivity between Prefrontal Cortex and Striatum during Category Learning. Neuron 83, 216–225 (2014).

21. M Abeles, Corticonics: Neural Circuits of the Cerebral Cortex. (Cambridge University Press, Cambridge), (1991).

22. D Fitzpatrick, JS Lund, D Schmechel, AC Towles, Distribution of GABAergic neurons and axon terminals in the macaque striate cortex. The J. Comp. Neurol. 264, 73–91 (1987).

23. SH Hendry, HD Schwark, EG Jones, J Yan, Numbers and proportions of GABA-immunoreactive neurons in different areas of monkey cerebral cortex. The J. Neurosci. The Off. J. Soc. for Neurosci. 7, 1503–1519 (1987).

24. R Coultrip, R Granger, G Lynch, A cortical model of winner-take-all competition via lateral inhibition. Neural Networks 5, 47–54 (1992).

25. ZH Mao, SG Massaquoi, Dynamics of Winner-Take-All Competition in Recurrent Neural Networks With Lateral Inhibition. IEEE Transactions on Neural Networks 18, 55–69 (2007).

26. S Kaski, T Kohonen, Winner-take-all networks for physiological models of competitive learning. Neural Networks 7, 973–984 (1994).

27. SI Amari, A Mathematical Approach to Neural Systems in Systems Neuroscience, ed. J Metzler. (Academic Press), pp. 67–117 (1977).

28. DJ Pinto, GB Ermentrout, Spatially Structured Activity in Synaptically Coupled Neuronal Networks: II. Lateral Inhibition and Standing Pulses. SIAM J. on Appl. Math. 62, 226–243 (2001).

29. E Oja, Neural networks, principal components, and subspaces. Int. J. Neural Syst. 01, 61–68 (1989).

30. S Grossberg, Competitive learning: From interactive activation to adaptive resonance. Cogn. Sci. 11, 23–63 (1987).

31. C von der Malsburg, Self-organization of orientation sensitive cells in the striate cortex. Kybernetik 14, 85–100 (1973).

32. S Coombes, A Byrne, Next Generation Neural Mass Models in Nonlinear Dynamics in Computational Neuroscience, eds. F Corinto, A Torcini. (Springer International Publishing, Cham), pp. 1–16 (2019).

33. A Byrne, RD O’Dea, M Forrester, J Ross, S Coombes, Next-generation neural mass and field modeling. J. Neurophysiol. 123, 726–742 (2020).

34. JB Ding, JN Guzman, JD Peterson, JA Goldberg, DJ Surmeier, Thalamic Gating of Corticostriatal Signaling by Cholinergic Interneurons. Neuron 67, 294–307 (2010).

35. R Granger, Engines of the Brain: The Computational Instruction Set of Human Cognition. AI Mag. 27, 15–32 (2006).

36. S Bhattacharya, SL Brincat, M Lundqvist, EK Miller, Traveling waves in the prefrontal cortex during working memory. PLOS Comput. Biol. 18, e1009827 (2022).

37. ZW Davis, et al., Spontaneous traveling waves naturally emerge from horizontal fiber time delays and travel through locally asynchronous-irregular states. Nat. Commun. 12, 6057 (2021).

38. EP Stephen, et al., Broadband slow-wave modulation in posterior and anterior cortex tracks distinct states of propofol-induced unconsciousness. Sci. Reports 10, 13701 (2020).

39. EV Lubenov, AG Siapas, Hippocampal theta oscillations are travelling waves. Nature 459, 534–539 (2009).

40. P Dayan, LF Abbott, Theoretical Neuroscience: Computational and Mathematical Modeling of Neural Systems. (MIT Press), (2005) Google-Books-ID: fLT4DwAAQBAJ.

41. W Gerstner, WM Kistler, R Naud, L Paninski, Neuronal Dynamics: From Single Neurons to Networks and Models of Cognition. (Cambridge University Press, Cambridge), (2014).

42. Gq Bi, Mm Poo, Synaptic Modifications in Cultured Hippocampal Neurons: Dependence on Spike Timing, Synaptic Strength, and Postsynaptic Cell Type. J. Neurosci. 18, 10464–10472 (1998).

43. S Song, KD Miller, LF Abbott, Competitive Hebbian learning through spike-timing-dependent synaptic plasticity. Nat. Neurosci. 3, 919–926 (2000).

44. Y LeCun, Y Bengio, G Hinton, Deep learning. Nature 521, 436–444 (2015).

45. JL Elman, Finding Structure in Time. Cogn. Sci. 14, 179–211 (1990).

46. DC Knill, A Pouget, The Bayesian brain: the role of uncertainty in neural coding and computation. Trends Neurosci. 27, 712–719 (2004).

47. RS Sutton, AG Barto, Reinforcement Learning: An Introduction. (pnMIT Press Cambridge, USA), 1st edition, (1998).

48. JJ Hopfield, Neural networks and physical systems with emergent collective computational abilities. Proc. Natl. Acad. Sci. United States Am. 79, 2554–2558 (1982).

49. S Wu, Si Amari, H Nakahara, Population Coding and Decoding in a Neural Field: A Computational Study. Neural Comput. 14, 999–1026 (2002).

50. DE Rumelhart, GE Hinton, RJ Williams, Learning representations by back-propagating errors. Nature 323, 533–536 (1986).

51. R Milo, et al., Network Motifs: Simple Building Blocks of Complex Networks. Science 298, 824–827 (2002).

52. O Sporns, R Kötter, Motifs in Brain Networks. PLOS Biol. 2, e369 (2004).

53. AP Georgopoulos, AB Schwartz, RE Kettner, Neuronal Population Coding of Movement Direction. Science 233, 1416–1419 (1986).

54. A Schurger, JD Sitt, S Dehaene, An accumulator model for spontaneous neural activity prior to self-initiated movement. Proc. Natl. Acad. Sci. 109, E2904–E2913 (2012).

55. B Libet, CA Gleason, EW Wright, DK Pearl, Time of conscious intention to act in relation to onset of cerebral activity (readiness-potential): the unconscious initiation of a freely voluntary act. Brain 106, 623–642 (1983).

56. CS Soon, M Brass, HJ Heinze, JD Haynes, Unconscious determinants of free decisions in the human brain. Nat. Neurosci. 11, 543–545 (2008).

57. JD Haynes, et al., Reading Hidden Intentions in the Human Brain. Curr. Biol. 17, 323–328 (2007).

58. H Jeong, et al., Mesolimbic dopamine release conveys causal associations. Science 378, eabq6740 (2022).

59. AL Hodgkin, AF Huxley, A quantitative description of membrane current and its application to conduction and excitation in nerve. The J. Physiol. 117, 500–544 (1952).

60. SR Wicks, CJ Roehrig, CH Rankin, A Dynamic Network Simulation of the Nematode Tap Withdrawal Circuit: Predictions Concerning Synaptic Function Using Behavioral Criteria. The J. Neurosci. 16, 4017–4031 (1996).

61. BE Jones, Arousal systems. Front. Biosci. 8, 438–451 (2003).

62. AC Felch, RH Granger, The hypergeometric connectivity hypothesis: Divergent performance of brain circuits with different synaptic connectivity distributions. Brain Res. 1202, 3–13 (2008).

63. Y Shimo, O Hikosaka, Role of tonically active neurons in primate caudate in reward-oriented saccadic eye movement. The J. Neurosci. 21, 7804–7814 (2001).

64. A Singh, SM Papa, Striatal oscillations in parkinsonian non-human primates. Neuroscience 449, 116–122 (2020).

65. M Deffains, et al., Subthalamic, not striatal, activity correlates with basal ganglia downstream activity in normal and parkinsonian monkeys. eLife 5, e16443 (2016).

66. EM Vazey, DE Moorman, G Aston-Jones, Phasic locus coeruleus activity regulates cortical encoding of salience information. Proc. Natl. Acad. Sci. 115, E9439–E9448 (2018).

67. AC Kreitzer, RC Malenka, Striatal plasticity and basal ganglia circuit function. Neuron 60, 543–554 (2008).

68. JNJ Reynolds, JR Wickens, Dopamine-dependent plasticity of corticostriatal synapses. Neural Networks 15, 507–521 (2002).

69. J Larson, P Xiao, G Lynch, Reversal of LTP by theta frequency stimulation. Brain Res. 600, 97–102 (1993).

70. J Larson, E Munkacsy, Theta-burst LTP. Brain Res. 1621, 38–50 (2015).

71. R Granger, J Whitson, J Larson, G Lynch, Non-Hebbian properties of long-term potentiation enable high-capacity encoding of temporal sequences. Proc. Natl. Acad. Sci. 91, 10104–10108 (1994).

72. MA Castro-Alamancos, BW Connors, Cellular Mechanisms of the Augmenting Response: Short-Term Plasticity in a Thalamocortical Pathway. The J. Neurosci. 16, 7742–7756 (1996).

73. MA Castro-Alamancos, BW Connors, Short-Term Plasticity of a Thalamocortical Pathway Dynamically Modulated by Behavioral State. Science 272, 274–277 (1996).

74. G Mongillo, O Barak, M Tsodyks, Synaptic Theory of Working Memory. Science 319, 1543–1546 (2008).

75. H Kamiya, RS Zucker, Residual Ca2 + and short-term synaptic plasticity. Nature 371, 603–606 (1994).

76. K Ibata, Q Sun, GG Turrigiano, Rapid Synaptic Scaling Induced by Changes in Postsynaptic Firing. Neuron 57, 819–826 (2008).

77. F Bouchacourt, TJ Buschman, A flexible model of working memory. Neuron 103, 147–160.e8 (2019).

78. NT Carnevale, ML Hines, The NEURON Book. (Cambridge University Press), 1 edition, (2006).

79. D Goodman, Brian: a simulator for spiking neural networks in Python. Front. Neuroinformatics 2 2008).

80. MO Gewaltig, A Morrison, HE Plesser, NEST by Example: An Introduction to the Neural Simulation Tool NEST in Computational Systems Neurobiology, ed. N Le Novere. (Springer Netherlands, Dordrecht), pp. 533–558 (2012).

81. P Sanz-Leon, et al., The Virtual Brain: a simulator of primate brain network dynamics. Front. Neuroinformatics 7, 10 (2013).

## References

1. EG Antzoulatos, EK Miller, Increases in Functional Connectivity between Prefrontal Cortex and Striatum during Category Learning. Neuron 83, 216–225 (2014).

2. EG Antzoulatos, EK Miller, Differences between Neural Activity in Prefrontal Cortex and Striatum during Learning of Novel Abstract Categories. Neuron 71, 243–249 (2011).

3. R Coultrip, R Granger, G Lynch, A cortical model of winner-take-all competition via lateral inhibition. Neural Networks 5, 47–54 (1992).

4. M Abeles, Corticonics: Neural Circuits of the Cerebral Cortex. (Cambridge University Press, Cambridge), (1991).

5. D Fitzpatrick, JS Lund, DE Schmechel, AC Towles, Distribution of GABAergic neurons and axon terminals in the macaque striate cortex. The J. Comp. Neurol. 264, 73–91 (1987).

6. SH Hendry, HD Schwark, EG Jones, J Yan, Numbers and proportions of GABA-immunoreactive neurons in different areas of monkey cerebral cortex. The J. Neurosci. The Off. J. Soc. for Neurosci. 7, 1503–1519 (1987).

7. V Braitenberg, A Schuz, Cortex: statistics and geometry of neuronal connectivity. (Springer, Berlin, Heidelberg), (1998).

8. J Szentagothai, The ‘module-concept’ in cerebral cortex architecture. Brain Res. 95, 475–496 (1975).

9. DH Hubel, TN Wiesel, S LeVay, HB Barlow, RM Gaze, Plasticity of ocular dominance columns in monkey striate cortex. Philos. Transactions Royal Soc. London. B, Biol. Sci. 278, 377–409 (1997).

10. OD Creutzfeldt, HC Nothdurft, Representation of complex visual stimuli in the brain. Naturwissenschaften 65, 307–318 (1978).

11. VB Mountcastle, Brain mechanisms for directed attention. J. Royal Soc. Medicine 71, 14–28 (1978).

12. A Keller, EL White, Triads: a synaptic network component in the cerebral cortex. Brain Res. 496, 105–112 (1989).

13. RAW Galuske, W Schlote, H Bratzke, W Singer, Interhemispheric asymmetries of the modular structure in human temporal cortex. Science 289, 1946–1949 (2000).

14. MS Gazzaniga, Regional differences in cortical organization. Science 289, 1887–1888 (2000).

15. MA Castro-alamancos, BW Connors, Thalamocortical synapses. Prog. Neurobiol. 51, 581–606 (1997).

16. EG Jones, Viewpoint: the core and matrix of thalamic organization. Neuroscience 85, 331–345 (1998).

17. AJ Heynen, MF Bear, Long-term potentiation of thalamocortical transmission in the adult visual cortex in vivo. J. Neurosci. 21, 9801–9813 (2001).

18. G Silberberg, A Gupta, H Markram, Stereotypy in neocortical microcircuits. Trends Neurosci. 25, 227–230 (2002).

19. F Valverde, Structure of the cerebral cortex. Intrinsic organization and comparative analysis of the neocortex. Revista de neurologia 34, 758–780 (2002).

20. EL White, A Peters, Cortical modules in the posteromedial barrel subfield (Sml) of the mouse. J. Comp. Neurol. 334, 86–96 (1993).

21. A Peters, BR Payne, J Budd, A numerical analysis of the geniculocortical input to striate cortex in the monkey. Cereb. Cortex 4, 215–229 (1994).

22. M Molinari, et al., Auditory thalamocortical pathways defined in monkeys by calcium-binding protein immunoreactivity. J. Comp. Neurol. 362, 171–194 (1995).

23. H Killackey, FF Ebner, Two different types of thalamocortical projections to a single cortical area in mammals. Brain Behav. Evol. 6, 156–169 (1972).

24. M Herkenham, New perspectives on the organization and evolution of nonspecific thalamocortical projections in Sensory-Motor Areas and Aspects of Cortical Connectivity, Cerebral Cortex, eds. EG Jones, A Peters. (Springer US, Boston, MA), pp. 403–445 (1986).

25. HJ Groenewegen, HW Berendse, JG Wolters, AHM Lohman, The anatomical relationship of the prefrontal cortex with the striatopallidal system, the thalamus and the amygdala: evidence for a parallel organization in Progress in Brain Research, The Prefrontal Its Structure, Function and Cortex Pathology, eds. HBM Uylings, CG Van Eden, JPC De Bruin, MA Corner, MGP Feenstra. (Elsevier) Vol. 85, pp. 95–118 (1991).

26. J DeFelipe, EG Jones, Parvalbumin immunoreactivity reveals layer IV of monkey cerebral cortex as a mosaic of microzones of thalamic afferent terminations. Brain Res. 562, 39–47 (1991).

27. T Van Groen, JM Wyss, Projections from the anterodorsal and anteroveniral nucleus of the thalamus to the limbic cortex in the rat. J. Comp. Neurol. 358, 584–604 (1995).

28. TF Freund, KaC Martin, D Whitteridge, Innervation of cat visual areas 17 and 18 by physiologically identified X- and Y-type thalamic afferents. I. Arborization patterns and quantitative distribution of postsynaptic elements. J. Comp. Neurol. 242, 263–274 (1985).

29. H Barbas, N Rempel-Clower, Cortical structure predicts the pattern of corticocortical connections. Cereb. Cortex 7, 635–646 (1997).

30. KS Rockland, Non-uniformity of extrinsic connections and columnar organization. J. Neurocytol. 31, 247–253 (2002).

31. HA Swadlow, AG Gusev, T Bezdudnaya, Activation of a cortical column by a thalamocortical impulse. J. Neurosci. 22, 7766–7773 (2002).

32. DB Bender, Visual activation of neurons in the primate pulvinar depends on cortex but not colliculus. Brain Res. 279, 258–261 (1983).

33. ME Diamond, M Armstrong-James, MJ Budway, FF Ebner, Somatic sensory responses in the rostral sector of the posterior group (POm) and in the ventral posterior medial nucleus (VPM) of the rat thalamus: Dependence on the barrel field cortex. J. Comp. Neurol. 319, 66–84 (1992).

34. ME Diamond, M Armstrong-James, FF Ebner, Somatic sensory responses in the rostral sector of the posterior group (POm) and in the ventral posterior medial nucleus (VPM) of the rat thalamus. J. Comp. Neurol. 318, 462–476 (1992).

35. JJ Chrobak, G Buzsaki, Gamma oscillations in the entorhinal cortex of the freely behaving rat. J. Neurosci. 18, 388–398 (1998).

36. M Sarter, JP Bruno, The neglected constituent of the basal forebrain corticopetal projection system: GABAergic projections. Eur. J. Neurosci. 15, 1867–1873 (2002).

37. P Fries, JH Reynolds, AE Rorie, R Desimone, Modulation of oscillatory neuronal synchronization by selective visual attention. Science 291, 1560–1563 (2001).

38. A Rozov, N Burnashev, B Sakmann, E Neher, Transmitter release modulation by intracellular Ca2+ buffers in facilitating and depressing nerve terminals of pyramidal cells in layer 2/3 of the rat neocortex indicates a target cell-specific difference in presynaptic calcium dynamics. The J. Physiol. 531, 807–826 (2001).

39. A Knoblauch, G Palm, Scene segmentation by spike synchronization in reciprocally connected visual areas. I. Local effects of cortical feedback. Biol. Cybern. 87, 151–167 (2002).

40. B Pesaran, JS Pezaris, M Sahani, PP Mitra, RA Andersen, Temporal structure in neuronal activity during working memory in macaque parietal cortex. Nat. Neurosci. 5, 805–811 (2002).

41. A Rodriguez, J Whitson, R Granger, Derivation and Analysis of Basic Computational Operations of Thalamocortical Circuits. J. Cogn. Neurosci. 16, 856–877 (2004).

42. J Ambros-Ingerson, R Granger, G Lynch, Simulation of paleocortex performs hierarchical clustering. Science 247, 1344–1348 (1990).

43. R Granger, J Whitson, J Larson, G Lynch, Non-Hebbian properties of long-term potentiation enable high-capacity encoding of temporal sequences. Proc. Natl. Acad. Sci. 91, 10104–10108 (1994).

44. A Bocincova, TJ Buschman, MG Stokes, SG Manohar, Neural signature of flexible coding in prefrontal cortex. Proc. Natl. Acad. Sci. 119, e2200400119 (2022).

45. R Granger, Brain Circuit Implementation: High-Precision Computation from Low-Precision Components in Toward Replacement Parts for the Brain, eds. T Berger, DL Glanzman. (The MIT Press), pp. 277–294 (2005).

46. R Granger, Engines of the Brain: The Computational Instruction Set of Human Cognition. AI Mag. 27, 15–32 (2006).

47. JJ Canales, et al., Shifts in striatal responsivity evoked by chronic stimulation of dopamine and glutamate systems. Brain 125, 2353–2363 (2002).

48. GE Alexander, MR DeLong, PL Strick, Parallel organization of functionally segregated circuits linking basal ganglia and cortex. Annu. Rev. Neurosci. 9, 357–381 (1986).

49. RL Reep, JV Corwin, Topographic organization of the striatal and thalamic connections of rat medial agranular cortex. Brain Res. 841, 43–52 (1999).

50. J Feingold, DJ Gibson, B DePasquale, AM Graybiel, Bursts of beta oscillation differentiate postperformance activity in the striatum and motor cortex of monkeys performing movement tasks. Proc. Natl. Acad. Sci. 112, 13687–13692 (2015).

51. M Lundqvist, et al., Gamma and Beta Bursts Underlie Working Memory. Neuron 90, 152–164 (2016).

52. M Lundqvist, et al., Working memory control dynamics follow principles of spatial computing. Nat. Commun. 14, 1429 (2023).

53. A Reiner, N Hart, W Lei, Y Deng, Corticostriatal projection neurons-dichotomous types and dichotomous functions. Front. Neuroanat. 4 (2010).

54. AM Graybiel, T Aosaki, AW Flaherty, M Kimura, The basal ganglia and adaptive motor control. Science 265, 1826–1831 (1994).

55. BJ Hunnicutt, et al., A comprehensive excitatory input map of the striatum reveals novel functional organization. eLife 5, e19103 (2016).

56. W Schultz, P Dayan, PR Montague, A Neural Substrate of Prediction and Reward. Science 275, 1593–1599 (1997).

57. W Schultz, Predictive reward signal of dopamine neurons. J. Neurophysiol. 80, 1–27 (1998).

58. W Schultz, Updating dopamine reward signals. Curr. Opin. Neurobiol. 23, 229–238 (2013).

59. W Schultz, Dopamine reward prediction-error signalling: a two-component response. Nat. Rev. Neurosci. 17, 183–195 (2016).

60. BE Jones, Arousal systems. Front. Biosci. 8, 438–451 (2003).

61. RE Brown, JT McKenna, Turning a Negative into a Positive: Ascending GABAergic Control of Cortical Activation and Arousal. Front. Neurol. 6, 135 (2015).

62. TF Freund, V Meskenaite, gamma-Aminobutyric acid-containing basal forebrain neurons innervate inhibitory interneurons in the neocortex. Proc. Natl. Acad. Sci. 89, 738–742 (1992).

63. EM Vazey, DE Moorman, G Aston-Jones, Phasic locus coeruleus activity regulates cortical encoding of salience information. Proc. Natl. Acad. Sci. 115, E9439–E9448 (2018).

64. BE Kilavik, M Zaepffel, A Brovelli, WA MacKay, A Riehle, The ups and downs of beta oscillations in sensorimotor cortex. Exp. Neurol. 245, 15–26 (2013).

65. MA Sherman, et al., Neural mechanisms of transient neocortical beta rhythms: Converging evidence from humans, computational modeling, monkeys, and mice. Proc. Natl. Acad. Sci. 113, E4885–E4894 (2016).

66. Y Kawaguchi, T Shindou, Noradrenergic excitation and inhibition of GABAergic cell types in rat frontal cortex. J. Neurosci. 18, 6963–6976 (1998) Publisher: Society for Neuroscience Section: ARTICLE.

67. H Salgado, et al., Layer-specific noradrenergic modulation of inhibition in cortical layer II/III. Cereb. Cortex 21, 212–221 (2011).

68. AI Gulyas, N Hajos, TF Freund, Interneurons containing calretinin are specialized to control other interneurons in the rat hippocampus. J. Neurosci. 16, 3397–3411 (1996).

69. JM Blasco-Ibanez, FJ Martinez-Guijarro, TF Freund, Enkephalin-containing interneurons are specialized to innervate other interneurons in the hippocampal CA1 region of the rat and guinea-pig. Eur. J. Neurosci. 10, 1784–1795 (1998).

70. AI Gulyas, L Acsady, TF Freund, Structural basis of the cholinergic and serotonergic modulation of GABAergic neurons in the hippocampus. Neurochem. Int. 34, 359–372 (1999).

71. TW Kim, et al., Selective localization of amyloid precursor-like protein 1 in the cerebral cortex postsynaptic density. Mol. Brain Res. 32, 36–44 (1995).

72. JJ Zhu, BW Connors, Intrinsic firing patterns and whisker-evoked synaptic responses of neurons in the rat barrel cortex. J. Neurophysiol. 81, 1171–1183 (1999).

73. P Reinagel, D Godwin, SM Sherman, C Koch, Encoding of visual information by lgn bursts. J. Neurophysiol. 81, 2558–2569 (1999).

74. NN Odean, M Sanayei, MN Shadlen, Transient oscillations of neural firing rate associated with routing of evidence in a perceptual decision. J. Neurosci. 43, 6369–6383 (2023).

75. EA Kramar, B Lin, CS Rex, CM Gall, G Lynch, Integrin-driven actin polymerization consolidates long-term potentiation. Proc. Natl. Acad. Sci. 103, 5579–5584 (2006).

76. J Larson, P Xiao, G Lynch, Reversal of LTP by theta frequency stimulation. Brain Res. 600, 97–102 (1993).

77. J Larson, E Munkacsy, Theta-burst LTP. Brain Res. 1621, 38–50 (2015).

78. DD Mott, DV Lewis, Facilitation of the Induction of Long-term Potentiation by GABAB Receptors. Science 252, 1718–1720 (1991) Publisher: American Association for the Advancement of Science.

79. DD Mott, DV Lewis, GABAB receptors mediate disinhibition and facilitate long-term potentiation in the dentate gyrus. Epilepsy research Suppl. 7, 119–134 (1992).

80. T Nakashiba, et al., Young Dentate Granule Cells Mediate Pattern Separation, whereas Old Granule Cells Facilitate Pattern Completion. Cell 149, 188–201 (2012).

81. A Terashima, et al., An Essential Role for PICK1 in NMDA Receptor-Dependent Bidirectional Synaptic Plasticity. Neuron 57, 872–882 (2008).

82. N Gorelova, JK Seamans, CR Yang, Mechanisms of Dopamine Activation of Fast-Spiking Interneurons That Exert Inhibition in Rat Prefrontal Cortex. J. Neurophysiol. 88, 3150–3166 (2002).

83. P Zhong, Z Yan, Distinct Physiological Effects of Dopamine D4 Receptors on Prefrontal Cortical Pyramidal Neurons and Fast-Spiking Interneurons. Cereb. Cortex 26, 180–191 (2016).

84. A Kirkwood, SM Dudek, JT Gold, CD Aizenman, MF Bear, Common forms of synaptic plasticity in the hippocampus and neocortex in vitro. Science 260, 1518–1521 (1993).

85. MF Bear, A Kirkwood, Neocortical long-term potentiation. Curr. Opin. Neurobiol. 3, 197–202 (1993).

86. MA Castro-Alamancos, JP Donoghue, BW Connors, Different forms of synaptic plasticity in somatosensory and motor areas of the neocortex. J. Neurosci. 15, 5324–5333 (1995).

87. DV Buonomano, MM Merzenich, Cortical plasticity: from synapses to maps. Annu. Rev. Neurosci. 21, 149–186 (1998).

88. K Seki, M Kudoh, K Shibuki, Sequence dependence of post-tetanic potentiation after sequential heterosynaptic stimulation in the rat auditory cortex. The J. Physiol. 533, 503–518 (2001).

89. G Mongillo, O Barak, M Tsodyks, Synaptic Theory of Working Memory. Science 319, 1543–1546 (2008).

90. L Kozachkov, et al., Robust and brain-like working memory through short-term synaptic plasticity. PLOS Comput. Biol. 18, e1010776 (2022).

91. T Aosaki, et al., Responses of tonically active neurons in the primate’s striatum undergo systematic changes during behavioral sensorimotor conditioning. J. Neurosci. 14, 3969–3984 (1994).

92. T Aosaki, M Kimura, AM Graybiel, Temporal and spatial characteristics of tonically active neurons of the primate’s striatum. J. Neurophysiol. 73, 1234–1252 (1995).

93. A Raz, A Feingold, V Zelanskaya, E Vaadia, H Bergman, Neuronal synchronization of tonically active neurons in the striatum of normal and parkinsonian primates. J. Neurophysiol. 76, 2083–2088 (1996).

94. H Inokawa, N Matsumoto, M Kimura, H Yamada, Tonically active neurons in the monkey dorsal striatum signal outcome feedback during trial-and-error search behavior. Neuroscience 446, 271–284 (2020).

95. Y Shimo, O Hikosaka, Role of tonically active neurons in primate caudate in reward-oriented saccadic eye movement. The J. Neurosci. 21, 7804–7814 (2001).

96. YF Zhang, SJ Cragg, Pauses in Striatal Cholinergic Interneurons: What is Revealed by Their Common Themes and Variations? Front. Syst. Neurosci. 11 (2017).

97. AC Martel, et al., Targeted transgene expression in cholinergic interneurons in the monkey striatum using canine adenovirus serotype 2 vectors. Front. Mol. Neurosci. 13 (2020).

98. YD Van der Werf, MP Witter, HJ Groenewegen, The intralaminar and midline nuclei of the thalamus. Anatomical and functional evidence for participation in processes of arousal and awareness. Brain Res. Rev. 39, 107–140 (2002).

99. G Mandelbaum, et al., Distinct cortical-thalamic-striatal circuits through the parafascicular nucleus. Neuron 102, 636–652.e7 (2019).

100. BD Bennett, CJ Wilson, Spontaneous activity of neostriatal cholinergic interneurons in vitro. J. Neurosci. 19, 5586–5596 (1999).

101. O Hikosaka, M Sakamoto, S Usui, Functional properties of monkey caudate neurons. I. Activities related to saccadic eye movements. J. Neurophysiol. 61, 780–798 (1989).

102. P Apicella, Leading tonically active neurons of the striatum from reward detection to context recognition. Trends Neurosci. 30, 299–306 (2007).

103. N Abudukeyoumu, T Hernandez-Flores, M Garcia-Munoz, GW Arbuthnott, Cholinergic modulation of striatal microcircuits. Eur. J. Neurosci. 49, 604–622 (2019).

104. AL Hodgkin, AF Huxley, A quantitative description of membrane current and its application to conduction and excitation in nerve. The J. Physiol. 117, 500–544 (1952).

105. XJ Wang, Ionic basis for intrinsic 40 Hz neuronal oscillations:. NeuroReport 5, 221–224 (1993).

106. XJ Wang, Pacemaker Neurons for the Theta Rhythm and Their Synchronization in the Septohippocampal Reciprocal Loop. J. Neurophysiol. 87, 889–900 (2002).

107. C Koch, I Segev, TJ Sejnowski, TA Poggio, eds., Methods in Neuronal Modeling: From Ions to Networks, Computational Neuroscience Series. (A Bradford Book, Cambridge, MA, USA), 2 edition, (1998).

108. SR Wicks, CJ Roehrig, CH Rankin, A Dynamic Network Simulation of the Nematode Tap Withdrawal Circuit: Predictions Concerning Synaptic Function Using Behavioral Criteria. The J. Neurosci. 16, 4017–4031 (1996).

109. S Coombes, A Byrne, Next Generation Neural Mass Models in Nonlinear Dynamics in Computational Neuroscience, eds. F Corinto, A Torcini. (Springer International Publishing, Cham), pp. 1–16 (2019).

110. A Byrne, RD O’Dea, M Forrester, J Ross, S Coombes, Next-generation neural mass and field modeling. J. Neurophysiol. 123, 726–742 (2020).

111. GB Ermentrout, N Kopell, Parabolic Bursting in an Excitable System Coupled with a Slow Oscillation. SIAM J. on Appl. Math. 46, 233–253 (1986).

112. JB Ding, JN Guzman, JD Peterson, JA Goldberg, DJ Surmeier, Thalamic Gating of Corticostriatal Signaling by Cholinergic Interneurons. Neuron 67, 294–307 (2010).

113. MC Avery, JL Krichmar, Neuromodulatory systems and their interactions: a review of models, theories, and experiments. Front. Neural Circuits 11 (2017).

114. H Alitto, Y Dan, Cell-type-specific modulation of neocortical activity by basal forebrain input. Front. Syst. Neurosci. 6 (2013).

115. M Jayachandran, et al., Nucleus reuniens transiently synchronizes memory networks at beta frequencies. Nat. Commun. 14, 4326 (2023) Number: 1 Publisher: Nature Publishing Group.

116. A Bibbig, et al., Beta Rhythms (15–20 Hz) Generated by Nonreciprocal Communication in Hippocampus. J. Neurophysiol. 97, 2812–2823 (2007) Publisher: American Physiological Society.

117. LR Silva, Y Amitai, BW Connors, Intrinsic oscillations of neocortex generated by layer 5 pyramidal neurons. Sci. (New York, N.Y.) 251, 432–435 (1991).

118. Y Tachibana, H Iwamuro, H Kita, M Takada, A Nambu, Subthalamo-pallidal interactions underlying parkinsonian neuronal oscillations in the primate basal ganglia. Eur. J. Neurosci. 34, 1470–1484 (2011) _eprint: 10.1111/j.1460-9568.2011.07865.x.

119. MM McCarthy, et al., Striatal origin of the pathologic beta oscillations in Parkinson’s disease. Proc. Natl. Acad. Sci. 108, 11620–11625 (2011).

120. A Singh, SM Papa, Striatal oscillations in parkinsonian non-human primates. Neuroscience 449, 116–122 (2020).

121. V Breton-Provencher, M Sur, Active control of arousal by a locus coeruleus GABAergic circuit. Nat. Neurosci. 22, 218–228 (2019).

122. JY Hansen, et al., Integrating brainstem and cortical functional architectures (2023).

123. GMG Shepherd, Corticostriatal connectivity and its role in disease. Nat. Rev. Neurosci. 14, 278–291 (2013).

124. S Gee, et al., Synaptic activity unmasks dopamine D2 receptor modulation of a specific class of layer V pyramidal neurons in prefrontal cortex. J. Neurosci. 32, 4959–4971 (2012) Publisher: Society for Neuroscience Section: Articles.

125. HJ Seong, AG Carter, D1 receptor modulation of action potential firing in a subpopulation of layer 5 pyramidal neurons in the prefrontal cortex. J. Neurosci. 32, 10516–10521 (2012) Publisher: Society for Neuroscience Section: Brief Communications.

126. CR Gerfen, MN Economo, J Chandrashekar, Long distance projections of cortical pyramidal neurons. J. Neurosci. Res. 96, 1467–1475 (2018) _eprint: 10.1002/jnr.23978.

127. G Lynch, CS Rex, CM Gall, LTP consolidation: Substrates, explanatory power, and functional significance. Neuropharmacology 52, 12–23 (2007).

128. EO Neftci, BB Averbeck, Reinforcement learning in artificial and biological systems. Nat. Mach. Intell. 1, 133–143 (2019).

129. C Rackauckas, ODE Solver Multi-Language Wrapper Package Work-Precision Benchmarks (MATLAB, SciPy, Julia, deSolve (R)) (2023).

130. Y Ma, et al., ModelingToolkit: A Composable Graph Transformation System for Equation-Based Modeling. 2103.05244 (2021).

131. C Pantelides, The Consistent Initialization of Differential-Algebraic Systems. SIAM J. Scl. Stat. Comput. 9, 213–231 (1988).

132. K Shimako, M Koga, An Extension of Pantelides Algorithm for Consistent Initialization of Differential-Algebraic Equations Using Minimally Singularity. 2020 59th Annu. Conf. Soc. Instrum. Control. Eng. Jpn. (SICE) pp. 1676–1681 (2020).

133. M Otter, H Elmqvist, Transformation of Differential Algebraic Array Equations to Index One Form in Transformation of Differential Algebraic Array Equations to Index One Form. pp. 565–579 (2017).

134. H Elmqvist, T Henningsson, M Otter, Innovations for Future Modelica in Innovations for Future Modelica. pp. 693–702 (2017).

135. B Guenter, The D* Symbolic Differentiation Algorithm (2007).

136. AG Baydin, BA Pearlmutter, AA Radul, JM Siskind, Automatic Differentiation in Machine Learning: a Survey. 1502.05767 (2018).

137. C Rackauckas, N Qing, DifferentialEquations.jl - A Performant and Feature-Rich Ecosystem for Solving Differential Equations in Julia. J. Open Res. Softw. 5 (2017).

138. J Verner, Numerically optimal Runge–Kutta pairs with interpolants. Numer. Algorithms 53, 383–396 (2010).

139. C Torrence, GP Compo, A Practical Guide to Wavelet Analysis. Bull. Am. Meteorol. Soc. 79, 61–78 (1998) Publisher: American Meteorological Society Section: Bulletin of the American Meteorological Society.

